# RNA Helicase DDX5 Negatively Regulates Wnt Signaling and Hepatocyte Reprogramming in Hepatitis B Virus-related Hepatocellular Carcinoma

**DOI:** 10.1101/765388

**Authors:** Saravana Kumar Kailasam Mani, Zhibin Cui, Bingyu Yan, Sagar Utturkar, Adrien Foca, Nadim Fares, David Durantel, Nadia Lanman, Philippe Merle, Majid Kazemian, Ourania Andrisani

## Abstract

**Background and Aims:** RNA helicase DEAD box protein 5 (DDX5) is downregulated during hepatitis B virus (HBV) replication, and associates with poor prognosis HBV-related hepatocellular carcinoma (HCC). The aim of this study is to determine the mechanism and significance of DDX5 downregulation for HBV-driven HCC.

**Approach & Results:** We used established cellular models of HBV replication, HBV infection, as well as HBV-related liver tumors. HBV replicating *DDX5* knockdown hepatocytes were analyzed by RNAseq; differentially expressed genes were validated by qRT-PCR, and bioinformatic analyses of HCCs from The Cancer Genome Atlas. Our results show reduced expression of *DDX5* in HCCs of all etiologies is associated with poor survival. In HBV replicating hepatocytes, downregulation of DDX5 is mediated by miR17∼92 and miR106b∼25, induced by HBV infection. Increased expression of these miRNAs was quantified in HBV-associated HCCs expressing a hepatic cancer stem cell (hCSC)-like gene signature and reduced *DDX5* mRNA, suggesting a role for DDX5 in hCSC formation. Interestingly, DDX5 knockdown in HBV replicating hepatocyte cell lines resulted in hepatosphere formation, sorafenib and cisplatin resistance, Wnt signaling activation and pluripotency gene expression, all characteristics of hCSCs. Moreover, DDX5 knockdown increased viral replication. RNA-seq analyses of HBV-replicating DDX5 knockdown cells, identified enhanced expression of key genes of the Wnt/β-catenin pathway, including Frizzled7 (*FZD7)* and Matrix Metallopeptidase7 (*MMP7)*, indicative of Wnt signaling activation. Clinically, elevated *FZD7* expression correlates with poor patient survival. Importantly, inhibitors to miR17∼92 and miR106b∼25 restored DDX5 levels and suppressed both Wnt/β-catenin activation and viral replication.

**Conclusion:** DDX5 is a negative regulator of Wnt signaling and hepatocyte reprogramming in HCCs. Restoration of DDX5 levels in HBV-infected patients can exert both antitumor and antiviral effects.

## INTRODUCTION

Hepatocellular carcinoma (HCC) is a leading type of primary cancer with increasing incidence globally (1). Chronic Hepatitis B virus (HBV) infection remains one of the major etiologic factors in HCC pathogenesis (2). Despite the HBV vaccine, the WHO estimates globally 250 million people are chronically infected with HBV. Moreover, the HBV vaccine is not always protective. For instance, children born of infected mothers become chronically infected. Curative treatments for early stage disease include liver resection, transplantation, or local ablation. However, high recurrence rates after resection compromise patient outcomes. In advanced stage HCC, multi-kinase inhibitors including sorafenib (3), regorafenib (4), cabozantinib (5), lenvatinib or anti-angiogenic monoclonal antibodies such as ramucirumab (6) offer only palliative benefits. Thus, a compelling need exists to determine key molecular drivers of HCC pathogenesis, in order to design effective, targeted therapies that suppress both virus biosynthesis and liver cancer.

A cellular mechanism hijacked by HBV that regulates both host and viral gene transcription (7) involves the chromatin modifying Polycomb Repressive Complex 2 (PRC2) complex (8) and RNA helicase DDX5 (9). PRC2 represses transcription of genes by trimethylation of histone H3 on lysine 27 (H3K27me3). During HBV infection, viral covalently closed circular DNA (cccDNA) serving as template for viral transcription, assumes chromatin-like structure (10). Histone modifications associated with the HBV cccDNA/minichromosome determine viral transcription and replication rate (11). Our previous studies have shown that HBV replicating cells and HBV-related HCCs exhibit reduced PRC2 activity, resulting in de-repression of host PRC2 target genes (12–14). Moreover, knockdown of the essential PRC2 subunit SUZ12 enhances HBV replication (12, 13), suggesting loss of PRC2-mediated gene repression is advantageous for viral growth. Loss of PRC2 function occurs by proteasomal degradation of SUZ12, dependent on cellular polo-like-kinase1 (PLK1) (15), a host pro-viral factor (16). RNA helicases regulate a wide range of pathways (17). For example, DDX5 interacts with SUZ12, contributing to enhanced SUZ12 protein stability (7). Reduced DDX5 and SUZ12 protein levels correlate with enhanced viral transcription and replication, while in clinical samples, reduced *DDX5* expression correlates with hepatocyte de-differentiation, expression of PRC2 target genes (*EpCAM, NANOG, OCT4, SOX2*), and poor patient prognosis (7). These observations suggest a role for DDX5 in both HBV replication and HBV-induced HCC.

In this study, we investigated how HBV infection mediates DDX5 downregulation, and what are the consequences of DDX5 downregulation for the infected hepatocyte. We show that oncogenic miR-17∼92 and its paralog miR106b∼25 (18) directly target the three prime untranslated region (3’-UTR) of *DDX5*. miRNAs silence gene expression post-transcriptionally, regulating an array of biological processes, and linked to various diseased states including cancer (19). miR17∼92 is induced by c-Myc proto-oncogene (20), and miR106b∼25 is encoded within intron 13 of minichromosome maintenance complex component 7 (*MCM7)* (21). Importantly, both miRNAs are upregulated in HBV-induced HCCs (18, 22) and their over-expression is associated with liver cirrhosis and HCC (23), HBV replication, and HBV-associated HCC (18, 22). Biogenesis of mature miRNAs involves processing of primary miRNA (pri-miRNA) by the microprocessor complex (19). DDX5 is a critical component of this complex, and importantly, genes involved in the microprocessor complex are haploinsufficient tumor suppressors (24). Herein, we show DDX5 downregulation imparts cancer stem cell properties to hepatocytes, including hepatosphere formation, resistance to sorafenib and cisplatin, expression of pluripotency genes, and activation of Wnt signaling. Antagomirs (inhibitors) to these miRNAs restore DDX5 levels in HBV replicating cells, suppressing Wnt pathway activation and virus biosynthesis, i.e., acting as both antitumor and antiviral agents.

**Fig. 1.**
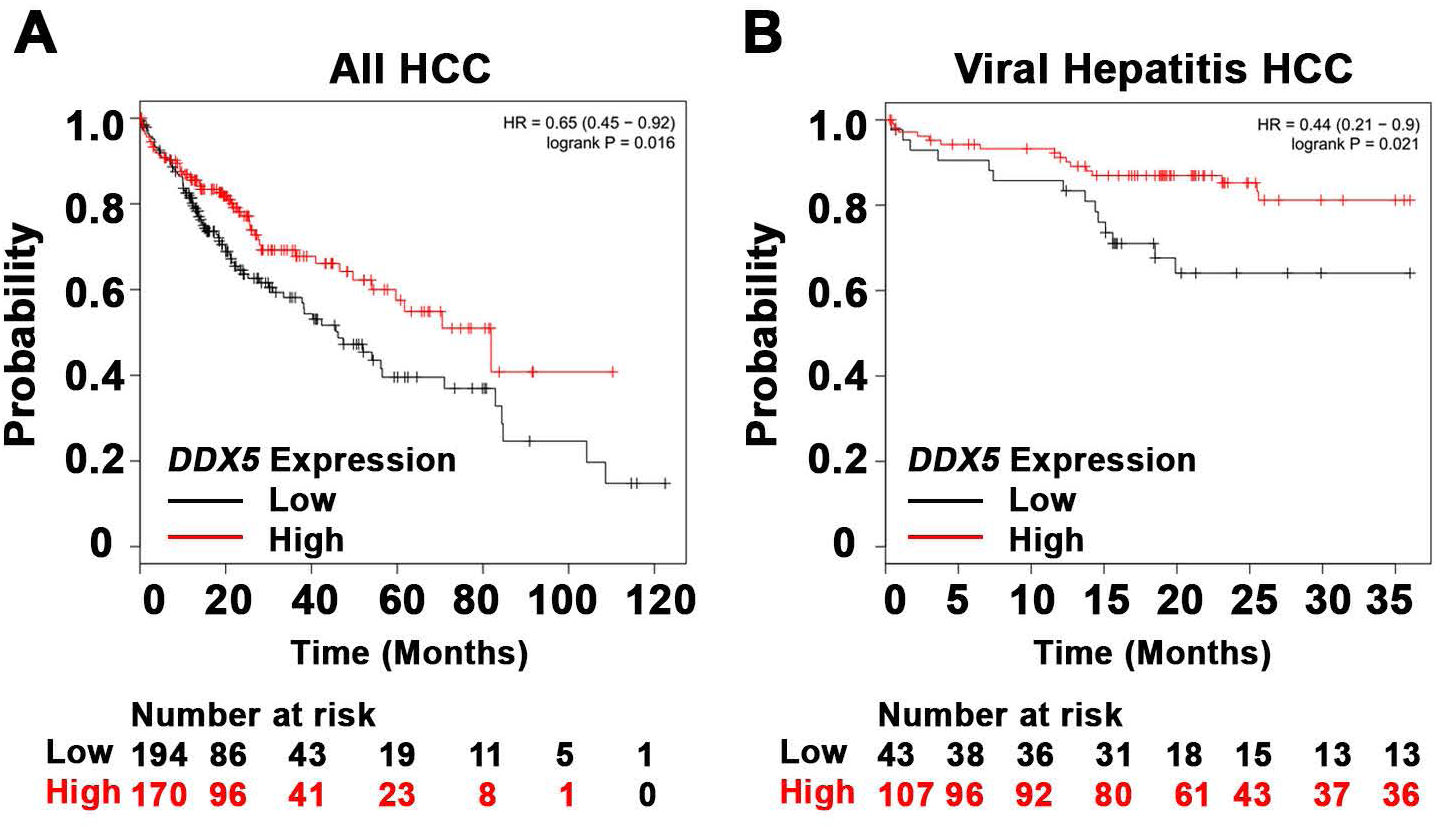
Reduced DDX5 levels in HCCs correlate with reduced patient survival. **(A-B)** Survival analysis of DDX5 expression in 364 HCC patients **(A)** and 150 hepatitis virus infected patients with HCC **(B)**. (The KMplotter database does not distinguish HCCs due to chronic HBV or HCV infection). Compared to patients with high DDX5 expression, patients with low DDX5 expression had statistically significant lower overall survival (OS) rate (P value<0.05). The best cutoff for DDX5 expression was auto selected by KMplotter.

## Materials and Methods

### Cell culture

Human HCC cell lines HepAD38 (25), HepG2 (7), HepG2-NTCP clone 7 (26) and HepaRG cells (27) were grown as described. Cell lines were routinely tested for mycoplasma. HepAD38 cell line and its derivatives were authenticated by short tandem repeat (STR) analysis performed by ATCC.

### Transfection and infection Assays

HepG2 and HepAD38 cells (5×10^4^ cells per well of 24 well-plate) were co-transfected with Renilla luciferase (25ng), Luc-3’UTR-*DDX5* (25ng), and control (Ctrl) vectors or plasmid encoding miR106b∼25 or miR17∼92, using Lipofectamine 3000 (Life Technologies). In HepAD38 cells (25), HBV replication was induced by tetracycline removal 48hrs prior to transfection. Luciferase activity was measured 48hrs after transfection using Dual Luciferase Assay system, per manufacturer’s protocol (Promega), and normalized to Renilla luciferase. Plasmids used are listed in Supporting Table S1. Infection assays of HepaRG and HepG2-NTCP cell lines were performed as described (26, 27), employing 100 HBV genome equivalents per cell.

### Wnt Reporter Assay

HBV replicating HepAD38 cells (5×10^4^ cells per well of 24 well-plate, day 3 of HBV replication) were co-transfected with TOPflash vector (25ng) containing TCF-binding sites upstream of firefly luciferase, and Renilla luciferase vector (25ng). Ctrl siRNA (40nM) or DDX5 siRNA (40nM) were co-transfected with Renilla and Firefly luciferase vectors using RNAiMax (Life Technologies). Luciferase activity was measured 48hrs after transfection using Dual Luciferase Assay system, per manufacturer’s protocol (Promega), and normalized to Renilla luciferase. Plasmids used are listed in **Table S1**.

### Sphere Assays

HBV replicating HepAD38 cells (1×10^3^) were seeded in ultra-low attachment 6-well plates (Corning). Cisplatin (10uM) and Sorafenib (2.5uM) were replaced every 3 days for 2 weeks, using sphere media containing DMEM/F12 (90% v/v), Penicillin/Streptomycin (1% v/v), G418 50 mg/ml (0.8% v/v), Fibroblast Growth factor 100 ng/μl (0.02% v/v), B27 (1X), and Epidermal growth factor 100 ng/μl (0.02% v/v).

### Cell Viability Assays

HBV replicating HepAD38 cells (1×10^4^) seeded in 96-well plates were treated with cisplatin (40 μM), sorafenib (7.5 μM), or DMSO for 24 hrs (day 5 of HBV replication). Growth inhibition measured at 490nm by CellTiter 96 AQ_ueous_ One Solution Cell Proliferation assay (Promega). 100 % viability refers to A_490_ value of DMSO-treated cells. Background absorbance measured from wells containing media and MTS without cells.

**Immunoblotting and Immunofluorescence microscopy** methods are described in detail in Supporting Information section. Antibodies employed are listed in **Table S2**.

**RNA preparation and qRT-PCR methods** are described in detail in in Supporting Information section; primer sequences are listed in **Table S3**, and reagents, chemical inhibitors and kits in **Table S4**.

### RNA-seq and Bioinformatics

HepAD38 cells, wild type (WT) and DDX5 knockdown (KD5), were grown +/− tetracycline for 10 days to induce HBV replication (25). Sorafenib (2.5 μM) treatment was for three days prior to harvesting. Three independent biological replicates were prepared for RNA isolation and RNA sequencing. Total RNA was submitted to Purdue Genomics Core Facility for quality assessment and next-generation sequencing. Paired-end 2×50 bp sequencing was performed using a HiSeq2500 system (Illumina). Sequencing data are available through the NCBI Gene Expression Omnibus (GEO) database (accession number GSE131257). Data quality control was performed using FastQC v0.11.8. The RNA expression level in each library was estimated by “rsem-calculate-expression” procedure in RSEM v1.3.112 using default parameters except “--bowtie-n 1 –bowtie-m 100 –seed-length 28 --paired-end”. The bowtie index required by RSEM software was generated by “rsem-prepare-reference” on all RefSeq genes, obtained from UCSC table browser on April 2017. EdgeR v3.24.013 package was used to normalize gene expression among all libraries and identify differentially expressed genes among samples with following constraints: fold change > 1.5, FDR < 0.05 and TPM > 1. Kaplan Meier (KM) survival plots were generated using Kaplan Meier Plotter (28). Gene set enrichment analysis (GSEA) was performed using GSEA software (29).

**Chromatin immunoprecipitation** (ChIP) assays were performed using Millipore ChIP Assay Kit (catalog no.: 17-295). Antibodies used are listed in **Table S2**. ChIP primers used are described in (20).

### Statistical Analysis

One way ANOVA with Sidak’s multiple comparison test with single pooled variance was performed using GraphPad Prism version 5.0 (GraphPad Software, San Diego, CA), comparing mean of each sample to mean of control (**Fig. 2B**, **Fig. 2C**, **Fig. 2E**). Two tailed t-test with Welch’s correction was used to determine significance in **Fig. 3E** and **Fig. 7B**. Two way ANOVA with Sidaks’s multiple comparison test was used comparing (i) mean of each microRNA in infected to uninfected samples (**Fig. 3A** and **Fig. 3B**); (ii) mean of each cell line to mean of WT cells (**Fig. 4D**, **Fig. 5A**, **Fig. 5C**); (iii) mean of each siRNA to control siRNA (**Fig. 5B**) (iv) mean of each inhibitor to mean of control inhibitor (**Fig. 8C**). Results were considered statistically significant if P<0.05.

**Fig. 2.**
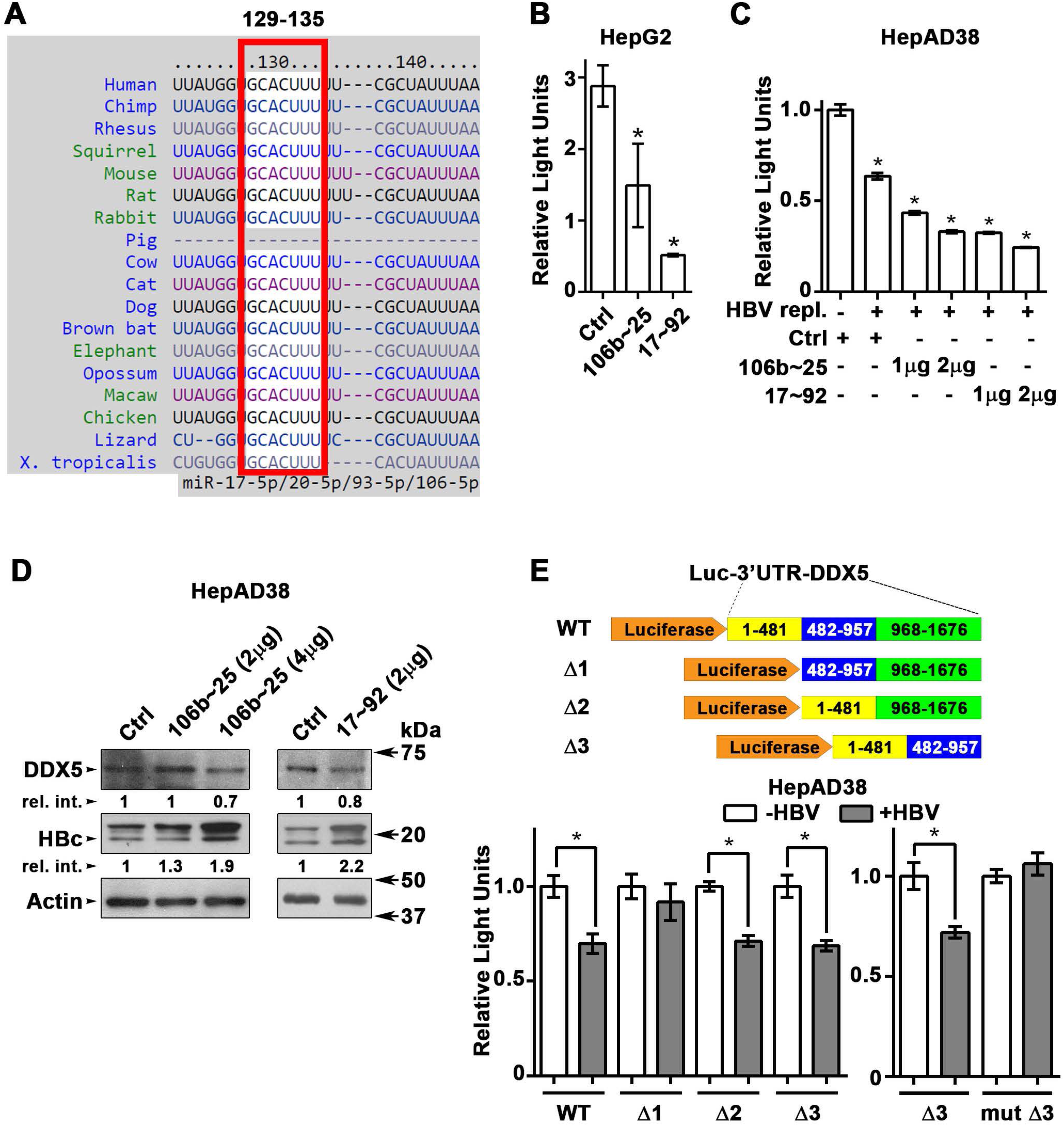
DDX5 is target of miR106b∼25 and miR17∼92. **(A)** Conserved seed sequences of miR17∼92 and miR106b∼25 in 3’UTR of DDX5, across 18 species. **(B-C)**, Transient transfections of Luc-3’UTR-DDX5 co-transfected with Renilla luciferase, and plasmids expressing mir106b∼25 or miR-17∼92 in HepG2 (**B)**, and HepAD38 (**C)** cells without (−) and with (+) HBV replication by tetracycline removal for 2 days. **(D)** Immunoblots of DDX5 and HBc following transfection of plasmids expressing mir106b∼25 or miR-17∼92, in HepAD38 cells with HBV replication for 2 days. Relative intensity (rel. int.) was quantified vs. actin using ImageJ software. (**E)** Luc-3’UTR-DDX5 containing the WT 3’UTR, indicated deletions Δ1, Δ2, Δ3 and site directed changes of nucleotides 129-135 (mut Δ3), co-transfected with Renilla luciferase in HepAD38 cells with (+) or without (−) HBV replication for 2 days (n=3). * p<0.05; Error bars indicate Mean ± SEM.

**Fig. 3.**
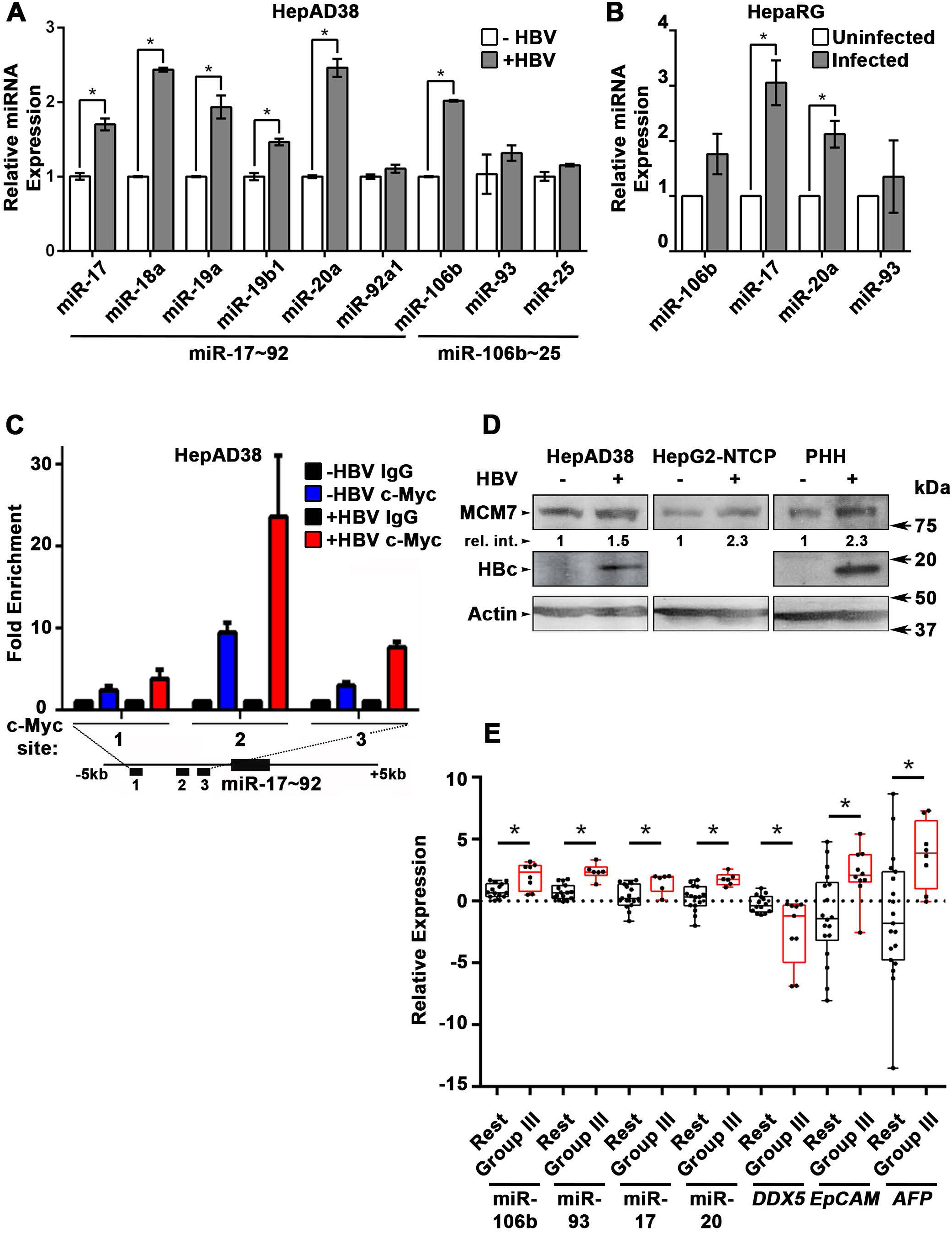
HBV replication induces expression of miR106b∼25 and miR17∼92. **(A)** qRT-PCR of indicated miRNAs in HepAD38 cells without (−) and with (+) HBV replication for 5 days (n=3 independent biological replicates). (**B**) qRT-PCR of indicated miRNAs in HBV-infected HepaRG cells on day4 post-infection (p.i.) (n=3 independent biological replicates). (**C)** ChIP assays in HepAD38 cells with (+) or without (−) HBV replication for 5 days, using c-Myc antibody and primer sets 1-3 spanning c-Myc binding sites (20). IgG was negative control. (n=3 independent biological replicates). Schematic representation of genomic interval encompassing the miR17∼92 cluster. RT-PCR amplicons are represented by numbered lines. (**D)** Immunoblots of MCM7 using lysates from HepAD38, HepG2-NTCP, and primary human hepatocytes (PHHs) with (+) or without (−) HBV replication/infection for 7days. (**E)** qRT-PCR of indicated miRNAs, and *DDX5* in HBV-related HCC tumor vs. peritumor samples. *EpCAM* and *AFP* (*α-fetoprotein*) comparison is between Tumor and Normal liver samples (14). Tumor samples were grouped according to (14) into Group III and Rest. * p<0.05; Error bars indicate Mean ± SEM.

**Fig. 4.**
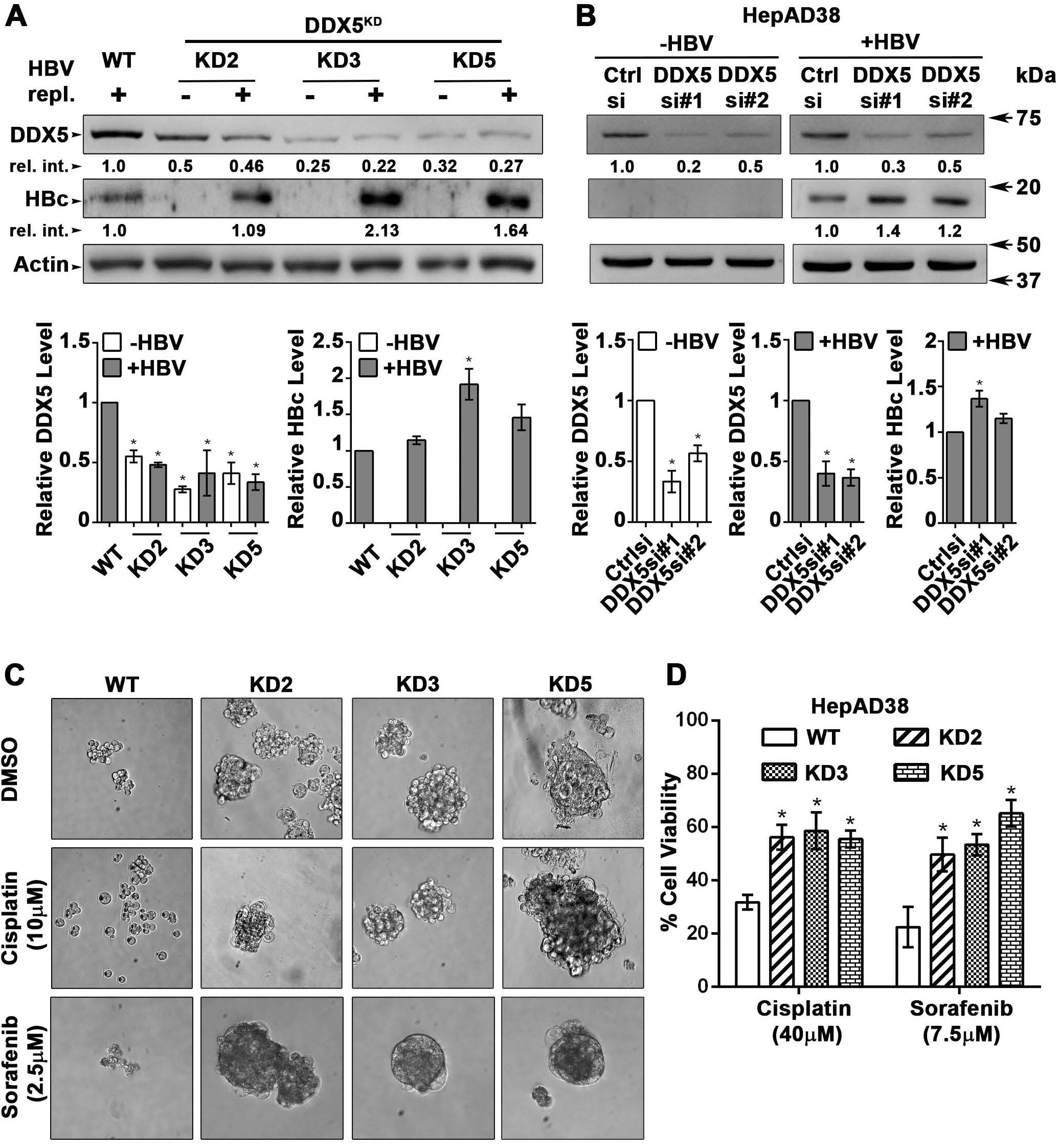
DDX5 knockdown confers stem cell-like properties. **(A-B)** Immunoblots of DDX5 and HBc in WT and DDX5 knockdown (KD2, KD3 and KD5) HepAD38 cells (**A**), and in WT HepAD38 cells (**B)** following transient transfection of two different siRNAs for DDX5 or control siRNA (Ctrl), without (−) and with (+) HBV replication for 5 days. Panels shown below the immunoblots are cumulative quantification of three independent biological replicates. (**C)** HBV replicating WT, KD2, KD3 and KD5 HepAD38 cells were grown for 14 days in hepatic sphere media using ultra-low attachment plates, with cisplatin (10 μM) or sorafenib (2.5 μM). Shown is a representative assay of three independent biological replicates. (**D)** Proliferation (MTS) assays performed with WT, KD2, KD3 and KD5 HepAD38 cells grown with (+) HBV replication for 5 days. Cells were treated with cisplatin (40μM), sorafenib (7.5 μM) or DMSO for 24 hours (n=3). *p<0.05; Error bars indicate Mean ± SEM.

**Fig. 5.**
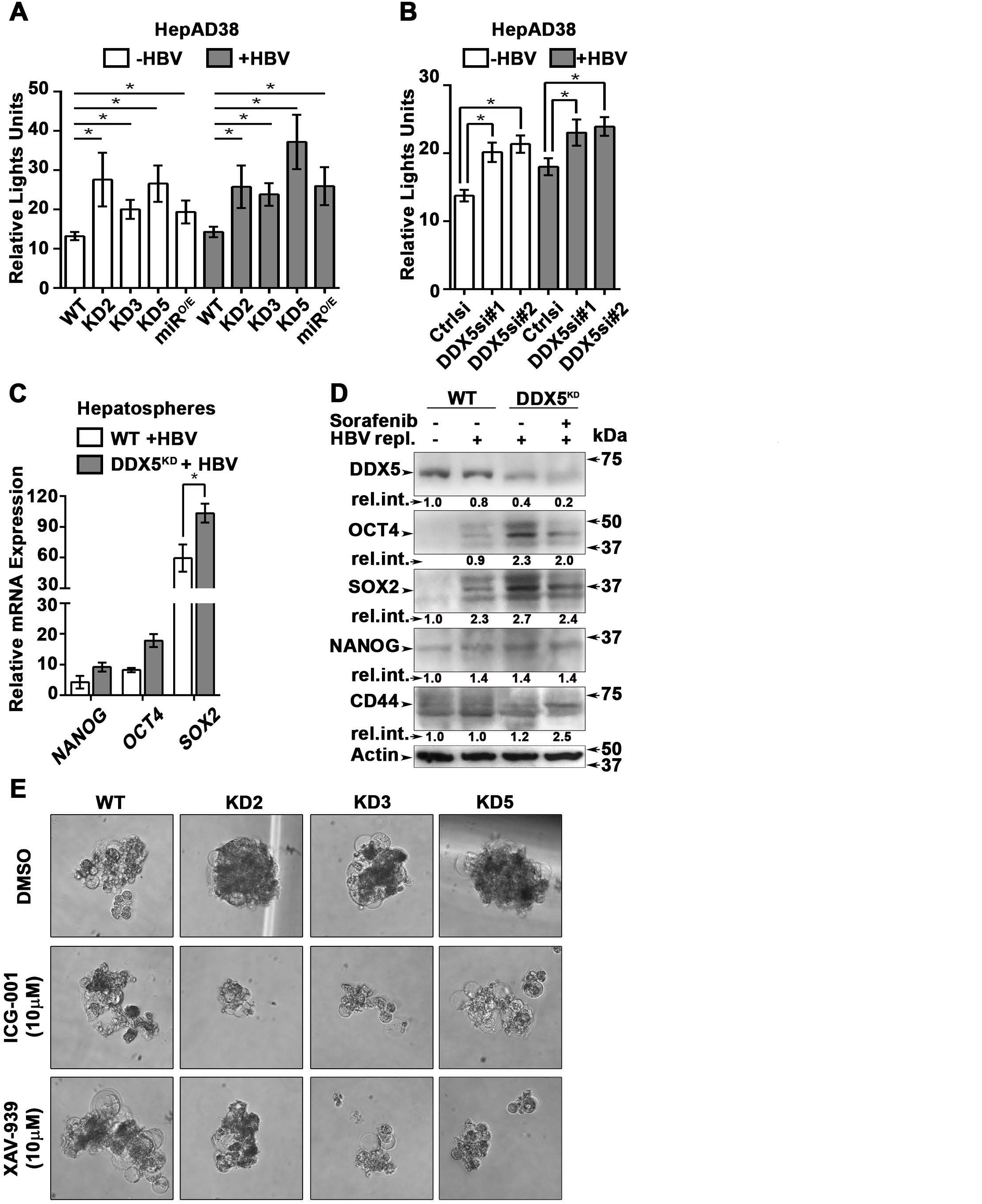
Downregulation of DDX5 activates Wnt signaling. (**A-B**) Transient co-transfections of TOPFlash and Renilla luciferase reporters in HepAD38 cells (WT, KD2, KD3, KD5 and miR^O/E^) **(A)** and HepAD38 cells transfected with *DDX5si#1, DDX5si#2* or negative control siRNA **(B)**. Luciferase activity from three independent assays, measured on day 5 of HBV replication are shown. (**C)** qRT-PCR of *OCT4 NANOG, SOX2* expression in WT and DDX5^KD^ HepAD38 hepatospheres with (+) HBV replication for 14 days. (**D)** Immunoblots of indicated proteins in WT and DDX5^KD^ HepAD38 hepatospheres with (+) HBV replication and sorafenib treatment for 14 days. **(E)** WT, KD2, KD3 and KD5 HepAD38 cells were grown for 14 days in hepatic sphere media using ultra-low attachment plates, in presence of Wnt inhibitors ICG-001 (10 μM) or XAV-939 (10 μM). Shown is a representative assay from three independent biological replicates. *p<0.05; Error bars indicate Mean ± SEM.

**Fig. 6.**
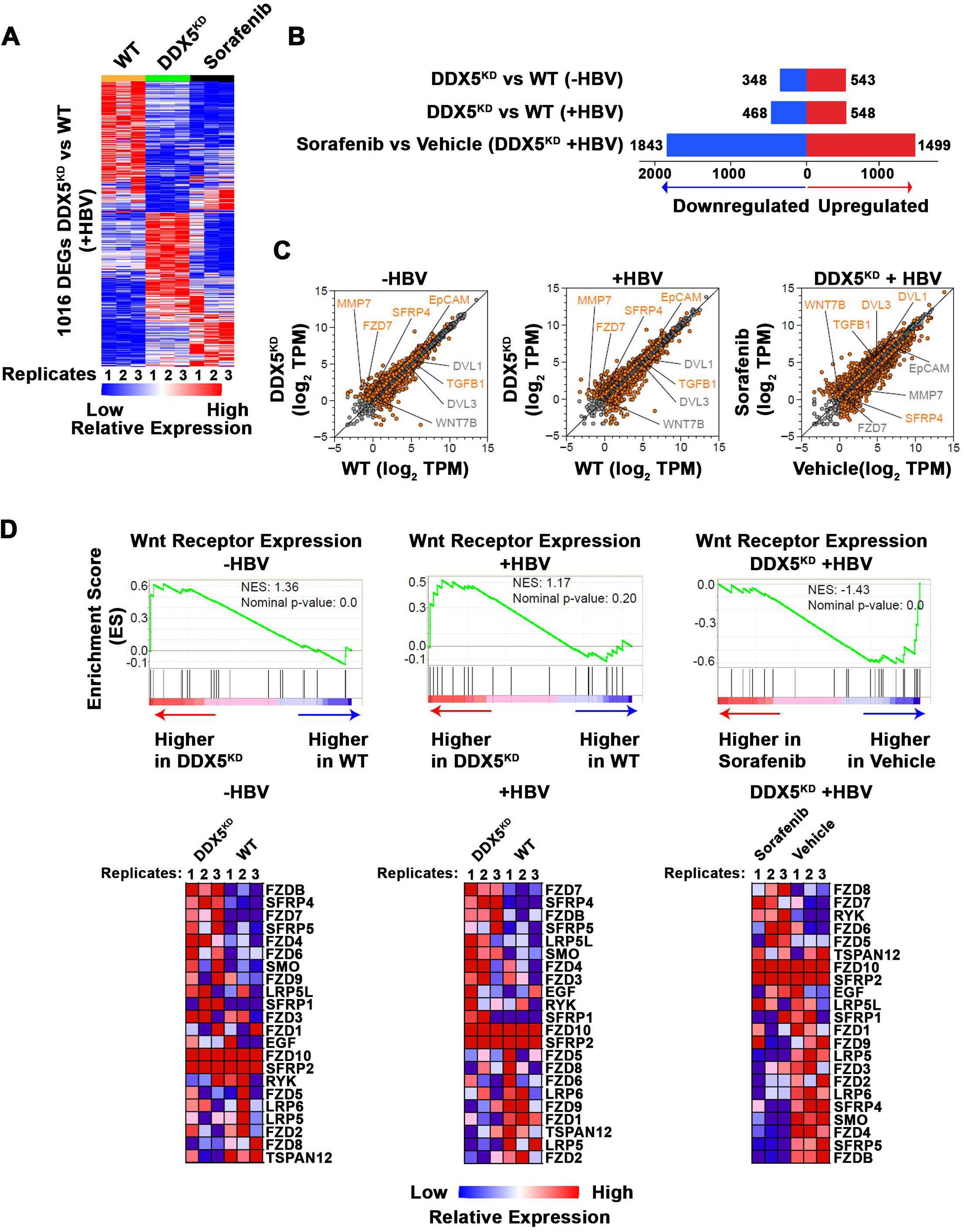
Transcriptomic analyses define dysregulated Wnt signaling in DDX5 knockdown cells. **(A)** Heat map of differentially expressed genes between DDX5^KD^ vs. WT HBV replicating cells for 10 days. RNA-seq samples are from three independent biological replicates. **(B)** Differentially expressed genes in three indicated comparisons: DDX5^KD^ vs. WT cells in the absence of HBV replication (-HBV), DDX5^KD^ vs. WT cells in the presence of HBV replication (+HBV), and DDX5^KD^ HBV replicating cells with or without sorafenib treatment. (**C**) Scatter plot showing mean gene expression values (n=3) in three above mentioned comparisons. Differentially expressed genes are highlighted in orange (FDR<0.05) and grey (FDR>0.05) (**D**) GSEA for Wnt receptor expression, comparing DDX5^KD^ vs. WT cells in the absence of HBV replication (left panel), DDX5^KD^ vs. WT cells in the presence (+) of HBV replication (middle panel), and DDX5^KD^ HBV replicating cells with or without sorafenib treatment (right panel). Heat maps of Wnt receptor expression (**GO: 0042813**) gene set.

**Fig. 7.**
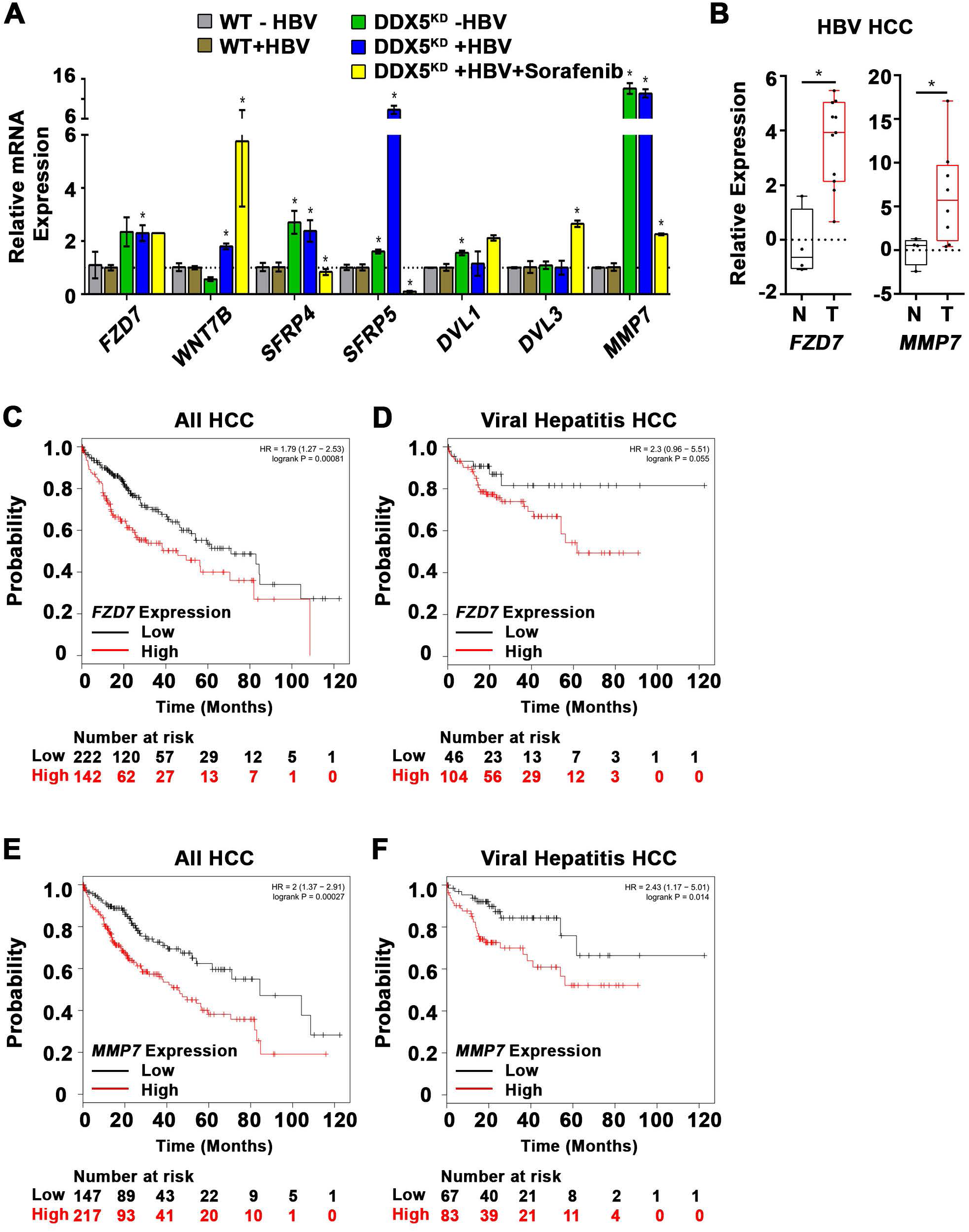
Genes upregulated by DDX5 knockdown are associated with poor overall survival in HCCs. **(A)** qRT-PCR validation of indicated genes using RNA isolated from HepAD38 cells, comparing DDX5^KD^ vs. WT cells in the absence (−) or presence (+) of HBV replication for 5 days, and with sorafenib (2.5 µM) treatment for 3 days. Expression values were calculated for WT -HBV vs. DDX5 -HBV; WT +HBV vs. DDX5 +HBV and DDX5+HBV vs. DDX5+HBV+Sorafenib, using ΔΔCt method. (n=3) **(B)** qRT-PCR of *FZD7* and *MMP7* employing RNA isolated from normal liver (N) vs. HBV-related HCCs (T) expressing reduced *DDX5* mRNA (7). RNA was isolated from the tumors listed in reference (7) and **Fig. S2**, namely tumors #13, #24, #52, #74, #76, #105, #141, #144, #150, #151. *p<0.05; Error bars indicate Mean ± SEM. **(C-D)** Survival analysis of *FZD7* expression in 364 HCC patients **(C)** or 150 viral hepatitis HCC patients **(D)**. **(E-F)** Survival analysis of *MMP7* expression in 364 HCC patients **(E)** or 150 viral hepatitis HCC patients **(F).**

**Fig. 8.**
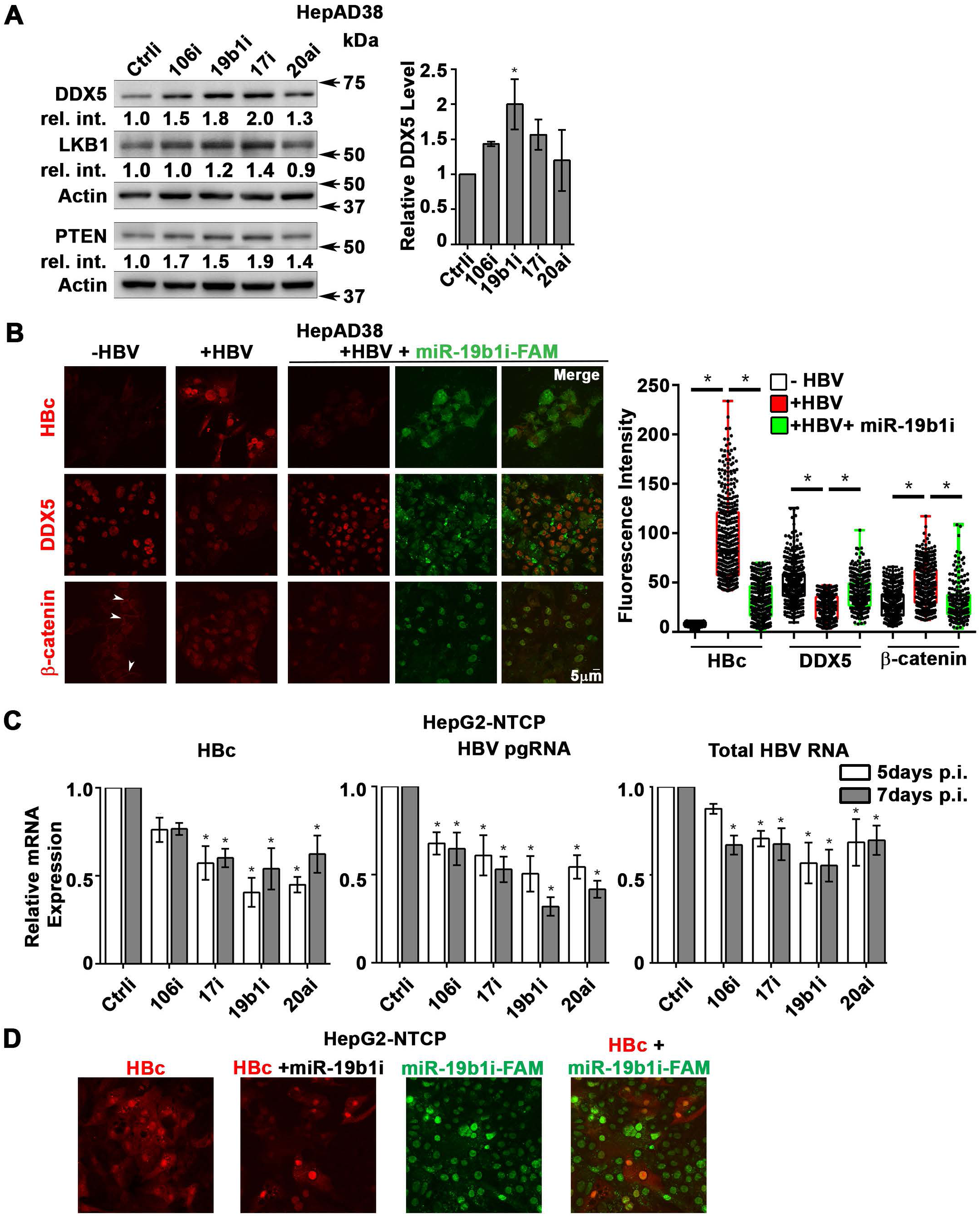
Antagomirs restore DDX5 in HBV replicating hepatocytes. **(A)** Immunoblots of DDX5, LKB1 and PTEN in HepAD38 cells with (+) HBV replication for 5 days. Cells were transfected for 24 hrs with 50 nM of indicated miRNA inhibitors/antagomirs (106i, 19b1i, 17i, 20ai) or control inhibitor (Ctrli), on day 4 of HBV replication. Cumulative immunoblot quantification of three independent biological replicates for DDX5 is shown on right panel, and for LKB1 and PTEN in **Fig. S6B**. *p<0.05; Error bars indicate Mean ± SEM. **(B**) Immunofluorescence confocal microscopy of HBc, DDX5 and β-catenin in HepAD38 cells without (−) or with (+) HBV replication for 5 days. Fluorescent antagomir for miR-19b1 (miR-19b1i-FAM, 50 nM) transfected on day 4 of HBV replication. White arrows indicate membrane localization of β-catenin. Shown is a representative assay from three independent biological replicates. Right panel is quantification of fluorescence intensities from 400 cells (mean gray value per μm^2^), employing ImageJ software. *p<0.05; Error bars indicate Mean ± SEM. **(C)** qRT-PCR of HBc RNA, HBV pgRNA and total HBV RNA levels in HepG2-NTCP cells infected with 100 HBV genome equivalents/cell. Infected cells were transfected with antagomirs of indicated miRNAs or Ctrli (50 nM) 24 hr prior to cell harvesting. Expression values calculated relative to Ctrli at 5 and 7 days p.i. using ΔΔCt method (n=3). *p<0.05; Error bars indicate Mean ± SEM. **(D**) Immunofluorescence confocal microscopy of HBc in HepG2-NTCP cells at 7 day p.i., infected with 100 HBV genome equivalents/cell. Antagomir miR-19b1i-FAM (50 nM) transfected on day 6 p.i. Shown is a representative assay from three independent biological replicates.

## RESULTS

### DDX5 downregulation by miR106b∼25 and miR17∼92

DDX5 is downregulated during HBV replication, and reduced levels of DDX5 in HBV-related HCCs tended towards poor patient prognosis (7). Here, we first established this link in HCCs of all etiologies (**Fig. 1A**), and in a large cohort of HCCs due to viral hepatitis (**Fig. 1B**), using Kaplan-Meier survival plots (28). Gender, ethnicity, pathological stages, and alcohol consumption did not seem to be confounders, as survival trends were poorer (although not always significant) in samples with lower DDX5 in any group (data not shown). We next studied DDX5 downregulation by miRNAs. Employing target prediction algorithm TargetScan, a highly conserved seed sequence for miR17∼92 and its paralog miR106b∼25 (18, 30) was identified in 3’ UTR of *DDX5*, located at nucleotides 129-135 (**Fig. 2A** and **Fig. S1**). To test whether miR106b∼25 and miR17∼92 downregulate DDX5, expression vectors encoding each of these miRNAs were co-transfected with expression vectors of Firefly luciferase gene fused to 3’UTR of *DDX5* (Luc-3’UTR-*DDX5*), and a Renilla luciferase vector for normalization of luciferase activity. Overexpression of either miR106b∼25 or miR17∼92 in HepG2 cells reduced Firefly luciferase activity, indicating a functional seed sequence in 3’UTR of *DDX5* (**Fig. 2B**). Likewise, transfection of Luc-3’UTR-*DDX5* vector in HepAD38 cells exhibited reduced luciferase activity upon induction of HBV replication, consistent with upregulation of both mi-RNAs in HBV-induced HCCs (18, 22) and their role in targeting DDX5. Overexpression of miR106b∼25 or miR17∼92 clusters further reduced luciferase activity (**Fig. 2C**). Importantly, overexpression of these miRNAs in HBV replicating HepAD38 cells (25) reduced protein level of endogenous DDX5, while levels of viral core antigen (HBc) were increased (**Fig. 2D**), suggesting loss of DDX5 is advantageous to viral biosynthesis. Finally, deletion analyses and site directed mutagenesis of 3’UTR of *DDX5* confirmed the presence of miRNA seed sequence at nucleotides 129-135 (**Fig. 2E**). Overall, our data indicate that *DDX5* mRNA is a direct target of miR106b∼25 and miR17∼92.

### HBV replication induces expression of miR106b∼25 and miR17∼92

Next, we quantified the expression of individual members of miR17∼92 and miR106b∼25 clusters in HBV replicating HepAD38 cells (**Fig. 3A**), and HBV infected HepaRG cells (**Fig. 3B**). Five out of six members of miR17∼92 cluster and one out of three members of miR106b∼25 cluster were significantly induced in HBV replicating HepAD38 cells (**Fig. 3A**), while two members of miR17∼92 were induced more than two-fold in HBV infected HepaRG cells (**Fig. 3B**). To understand how HBV infection increased miR17∼92 expression, we focused on miR17∼92 promoter region which contains three binding sites for c-Myc, often overexpressed in liver cancer and a key regulator of miR17∼92 (20). Two out of these three sites (i.e. sites #2 and #3) exhibited further increased c-Myc occupancy in HBV replicating HepAD38 cells (**Fig. 3C**), consistent with induction of miR17∼92 members (**Fig. 3A**). miR106b∼25 is encoded by intron13 of the *MCM7* gene and when *MCM7* is overexpressed, it also increases expression of miR106b∼25 (21). To determine whether *MCM7* is overexpressed, we quantified MCM7 protein levels in lysates from HBV replicating HepAD38 cells, HBV infected HepG2-NTCP cells, and HBV infected primary human hepatocytes (PHH). An approximately 2-fold increased MCM7 protein level was observed in all cases, supporting increased expression of miR106b∼25 in HBV replicating cells (**Fig. 3D**). Overall, these data indicate that HBV replication induces expression of miR106b∼25 and miR17∼92.

To determine whether miRNA-mediated downregulation of DDX5 is clinically relevant, expression of these miRNAs was quantified in HBV-related HCCs. Our previous study has identified a hepatic cancer stem cell (hCSC)-like gene signature associated with tumor recurrence after surgery (14). Interestingly, patients identified by this hCSC-like gene signature (denoted as Group III), exhibited increased expression of miR106b∼25 and miR17∼92, and reduced levels of *DDX5* mRNA (**Fig. 3E and Fig. S2**). Group III tumors also displayed statistically increased levels of *EpCAM*, a gene downstream of DDX5/PRC2 function (7), and *AFP*, a prognostic marker of HCC (**Fig. 3E**). These clinical data correlate high expression of miR106b∼25 and miR17∼92 to reduced *DDX5* levels, and in turn, tumor aggressiveness.

### DDX5 knockdown induces viral biosynthesis and confers cancer stem cell-like properties to hepatocytes

We next studied the functional significance of DDX5 loss to viral biosynthesis, and to hepatosphere formation. We derived three stable DDX5 knockdown cell lines (DDX5^KD^) from HepAD38 cells, referred to as KD2, KD3 and KD5 (**Fig. 4A**) using three different shRNAs. To assess whether DDX5 downregulation affected viral replication, we quantified HBc levels by immunoblots. HBc levels were increased after DDX5 knockdown, both in the stable cell lines (**Fig. 4A**) and after transient transfection of DDX5 siRNAs #1 and #2 (**Fig. 4B**). These results also agree with increased HBc levels observed upon overexpression of miR106b∼25 and miR17∼92 in HBV replicating HepAD38 cells (**Fig. 2D**). Collectively, these data suggest that DDX5 acts as a host restriction factor for HBV biosynthesis.

Since DDX5 was shown to act as a roadblock to pluripotency (31), we examined whether loss of DDX5 promotes a stem cell-like phenotype in HBV replicating hepatocytes. WT HepAD38 cells failed to form hepatospheres in ultra-low attachment plates. By contrast, DDX5^KD^ cells (KD2, KD3, and KD5) formed robust hepatospheres that survived treatment with chemotherapeutic drugs cisplatin (10μM) and sorafenib (2.5μM) (**Fig. 4C**). Furthermore, proliferation assays demonstrated reduced sensitivity to cisplatin and sorafenib upon DDX5 downregulation (**Fig. 4D** and **Fig. S3**), a characteristic feature of cancer stem cells (CSCs).

To determine pathways contributing to the observed stemness characteristics, we focused on Wnt/β-catenin signaling, one of the key pathways regulating stemness (32, 33). We performed luciferase assays to assess whether Wnt signaling was upregulated upon reduction of DDX5. Wnt-responsive TOPFlash reporter, containing LEF/TCF binding sites upstream of Firefly luciferase, was co-transfected with control Renilla luciferase into WT HepAD38 cells, HepAD38 cells overexpressing miR106b∼25 and miR17∼92 (miR^O/E^, Supporting **Fig. S4**), and stably or transiently DDX5 knockdown cells. Increased luciferase activity, i.e. increased Wnt/β-catenin pathway activation, was observed upon DDX5 downregulation by stable overexpression of miR106b∼25 and miR17∼92 (miR^O/E^), stable (KD2, KD3, and KD5) and transient (siRNAs #1 and #2) knockdown of DDX5 (**Fig. 5A** and **Fig. 5B**). Moreover, hepatospheres of HBV replicating DDX5 knockdown cells exhibited higher expression of pluripotency genes, determined by qRT-PCR (**Fig. 5C**) and immunoblotting of NANOG, SOX2, OCT4 and hCSC marker CD44 (**Fig. 5D**). Wnt inhibitors, ICG-001 (34) and XAV-939 (35), targeting different steps of Wnt signaling, suppressed hepatosphere formation, thereby linking Wnt pathway activation to the hCSC phenotype (**Fig. 5E**). Taken together, these data indicate that DDX5 maintains hepatocyte differentiation, and DDX5 downregulation promotes stemness via activation of Wnt signaling.

### Transcriptomic analyses define dysregulated Wnt signaling in DDX5 knockdown cells

To quantify the global effects of DDX5 knockdown on global gene expression in hepatocytes, we compared the transcriptome of WT vs. DDX5^KD^ HepAD38 cells as a function of HBV replication, and sorafenib treatment using RNA-seq. Three comparisons were performed, namely, DDX5^KD^ vs. WT cells in the absence of HBV replication (-HBV), DDX5^KD^ vs. WT cells in the presence of HBV replication (+HBV), and DDX5^KD^ HBV replicating cells with or without sorafenib treatment. Nearly 1000 genes were differentially expressed between DDX5^KD^ vs. WT cells (Fold change >1.5; FDR<0.05; **Fig. 6A-B**). Interestingly, sorafenib treatment exerted a highly pronounced effect on the number of both upregulated and downregulated genes (**Fig. 6B**). Importantly, DDX5 downregulation either in the absence or presence of HBV replication significantly increased *EpCAM* expression (**Fig. 6C**), which is a known hepatic stem cell marker and a Wnt/β-catenin signaling target gene (36). To further investigate the effect of DDX5 downregulation on Wnt/β-catenin signaling pathway, we asked whether expression of Wnt receptor genes differed between WT vs. DDX5^KD^ cells, using Gene Set Enrichment Analysis (GSEA)(29). We found that DDX5^KD^ cells have higher expression of several genes involved in Wnt receptor signaling, irrespective of HBV replication, consistent with activation of the Wnt pathway in DDX5^KD^ cells (**Fig. 5A and B**). For example, Wnt receptors (frizzled/*FZD*) were among the upregulated genes in DDX5^KD^ cells (**Fig. 6D**), in agreement with earlier studies linking Wnt receptor (*FZD7*) overexpression to Wnt pathway activation in HBV-related HCC (37). Interestingly, treatment of HBV replicating DDX5^KD^ cells with sorafenib reduced expression of genes which act as negative effectors of Wnt activation (38), specifically Secreted Frizzled Related Protein 4 (*SFRP4)* and *SFRP5* (**Fig. 6D**).

The expression of select genes involved in Wnt/β-catenin signaling was validated by qRT-PCR (**Fig. 7A**). Specifically, in DDX5^KD^ cells expression of *FZD7, MMP7, SFRP4* and *SFRP5* were increased irrespective of HBV replication (**Fig. 7A**). Sorafenib treatment of HBV replicating DDX5^KD^ cells increased expression of *WNT7B* ligand, while significantly suppressing expression of the negative regulators of Wnt signaling, *SFRP4* and *SFRP5* (**Fig. 7A**). To determine the clinical significance of our observations, we analyzed HBV-related HCC samples. Importantly, HBV-related HCCs with reduced *DDX5* expression exhibit increased *FZD7* and *MMP7* mRNA levels (**Fig. 7B**), further supporting the RNAseq data (**Fig. 6**). Clinically, increased expression levels of *FZD7* and *MMP7* is associated with poor overall survival of patients with HCC (**Fig. 7C and E**) and hepatitis virus-induced HCC (**Fig. 7D and F**). Likewise, reduced expression of *SFRP4* and enhanced expression of *DVL1, DVL3* (*39*) and *MMP7* exhibited poor overall survival even in sorafenib treated patients, suggesting a likely role of Wnt pathway activation in sorafenib resistance (Supporting **Fig. S5**). Taken together, our results demonstrate that downregulation of DDX5 is a key event leading to activation of Wnt signaling.

### miRNA inhibitors (antagomirs) restore DDX5 in HBV replicating hepatocytes

We have shown that DDX5 acts as a host cell restriction factor to HBV replication, a barrier to hepatocyte dedifferentiation, and a negative regulator of Wnt signaling (**Fig. 2D**, **Fig. 4A, B**). We have also shown that overexpression of miR106b∼25 and miR17∼92 downregulate DDX5 (**Fig. 2**). However, whether miRNA inhibitors can prevent DDX5 downregulation and associated phenotypes is unclear. To investigate the effect of the miRNA inhibitors (antagomirs), we targeted specific members of miR106b∼25 and miR17∼92 families. Tumor suppressors LKB1 (40), PTEN (21), a known negative regulator of activated phospho-AKT, and inhibitory SMAD7 (41) are well-described targets of miR17∼92 and miR106b∼25 clusters, respectively, and indeed, were downregulated in HBV replicating cells (supporting **Fig. S6A**). Transfection of indicated antagomirs restored DDX5 as well as tumor suppressors LKB1 and PTEN in HBV replicating cells (**Fig. 8A** and **Fig. S6B**), suggesting these antagomirs have antitumor potential.

Next, we analyzed the antagomir effect on HBV replication, employing immunofluorescence microscopy of HBc. Co-transfection of combination of antagomirs for miR-106b, miR-17, miR-20a, and miR-19b1 reduced HBc immunostaining by nearly 50% in HepAD38 cells (**Fig. S6C**). Accordingly, we designed antagomirs to miR-19b1 and miR-17 each conjugated to a fluorophore (FAM). HepAD38 cells were transfected with miR-19b1i-FAM (**Fig. 8B**) or miR-17i-FAM (**Fig. S6D)** on day3 of HBV replication. Inhibitor miR-19b1i-FAM reduced HBc immunofluorescence in HBV replicating cells, while the signal for DDX5 increased. Interestingly, miR-19b1i-FAM also reduced nuclear immunostaining of β-catenin in HBV replicating HepAD38 cells (**Fig. 8B**), indicating that restoring DDX5 suppressed Wnt/β-catenin pathway activation. Quantification of the fluorescence intensity demonstrates that the observed effects by miR-19b1i-FAM (**Fig. 8B**) and miR-17i-FAM (**Fig. S6D**) are statistically significant.

We then utilized the infection model of HepG2-NTCP cells to verify effects of miR106b∼25 and miR17∼92 antagomirs on virus biosynthesis. HepG2-NTCP cells were infected with 100 HBV genome equivalents per cell; on days 4 and 6 post infection (p.i.), HBV infected cells were transfected for 24hr with antagomirs for miR-106b, miR-17, miR-20a, and miR-19b1 or control inhibitor (Ctrli). Quantification of viral RNA on days 5 and 7 p.i. demonstrated significant reduction in the expression of HBc mRNA, pre-genomic RNA (pgRNA), and total HBV RNA in infected cells transfected with the indicated antagomirs (**Fig. 8C**). We also monitored HBV biosynthesis in HBV infected HepG2-NTCP cells by fluorescence microscopy for HBc, as a function of transfection of miR-19b1i-FAM (**Fig. 8D**) and miR-17i-FAM (**Fig. S6E**). On day 7 p.i., HBV infected cells with strong HBc immunofluorescence (red) exhibited low fluorescence due to miR-19b1i-FAM (green); conversely, cells with strong green fluorescence exhibited weak immunostaining for HBc (**Fig. 8D**). Similar results were obtained with miR-17i-FAM (**Fig. S6E**). Taken together, these results demonstrate that antagomirs for miR106b∼25 and miR17∼92 restore DDX5 levels in HBV infected cells, attenuating Wnt pathway activation and HBV replication.

## DISCUSSION

In this study, we investigated the mechanism by which HBV infection downregulates the RNA helicase DDX5, and the significance of this downregulation to HCC pathogenesis. RNA helicases are involved in all aspects of RNA metabolism, from transcription, epigenetic regulation, miRNA processing, to mRNA splicing, decay, and translation (9, 17). As a RNA helicase, DDX5 was shown to be a pro-viral host factor in biosynthesis of several RNA viruses, including HIV and HCV (42). By contrast, DDX5 has antiviral function in myxoma virus biosynthesis (43) and in HBV biosynthesis, by a mechanism not yet understood (7). In our earlier studies, we have observed that DDX5 protein levels become progressively reduced in HBV replicating/infected hepatocytes. In agreement with previous studies (7, 44), we also found that low DDX5 mRNA levels associate with poor prognosis of viral hepatitis-induced HCCs and HCCs of other etiologies (**Fig. 1**). Transcriptomic and functional analyses reveal that downregulation of DDX5 in HepAD38 hepatocytes results in activation of Wnt/β-catenin signaling (**Figs. 5** and **Fig. 6**), a pathway involved in reprogramming of hepatocytes towards a hCSC phenotype in HCCs (36, 45, 46). In addition, the Wnt/β-catenin pathway is an important regulator of adult liver size, liver regeneration, and metabolic zonation (47, 48). Whether DDX5 has a role in regulation of normal liver size, regeneration, and liver zonation is presently unknown.

In this study we show that downregulation of DDX5 during the course of HBV infection is mediated by miR106b∼25 (21) and miR17∼92 (20) (**Fig. 2**). These miRNAs were found upregulated during HBV replication (49) and in HBV-related HCCs (22). However, their role in HBV replication and virus-mediated hepatocarcinogenesis has been unknown. These miRNAs suppress pro-apoptotic functions of TGF-β pathway (30), and downregulate tumor suppressors PTEN (21) and LKB1 (40). Interestingly, as we demonstrate herein, HBV-induced miR106b∼25 and miR17∼92 target the same seed sequence in 3’UTR of *DDX5* and downregulate DDX5 during HBV replication (**Fig. 2**). The increased expression of *MCM7* encoding miR106b∼25 (21), and c-Myc-driven transcription of miR17∼92 (20) occurring during HBV replication offer mechanistic insights into the observed induction of these miRNAs (**Fig. 3**).

HBV-related HCCs display upregulated expression of these miRNAs (22). The highest induction of miR106b∼25 and miR17∼92 is observed in HBV-related HCCs that display the lowest *DDX5* mRNA level (**Fig. 3E**). Importantly, HCCs with low *DDX5* mRNA belong to Group III tumors that express the hCSC-like gene signature associated with poor patient prognosis (14). The expression levels of miR106b∼25, miR17∼92 and *DDX5* in these two groups of tumors, namely, Group III vs. the rest, display statistically significant differences (**Fig. 3E**), suggesting a link of DDX5 loss to a hCSC-like phenotype. Mechanistically, loss of function studies of DDX5 demonstrate that DDX5 is multifunctional, having a role in viral biosynthesis (**Fig. 4A** and **Fig. 4B**), and also in drug resistance and hepatocyte stemness (**Fig. 4C** and **Fig. 4D**). Specifically, HBV replicating DDX5^KD^ cells form hepatospheres when grown in low attachment plates, are resistant to cisplatin and sorafenib, and express elevated levels of pluripotency genes and hCSC marker CD44 (**Fig. 5**). Moreover, loss of DDX5 either by overexpression of miR106b∼25 and miR17∼92, stable knockdown of DDX5, or DDX5 siRNA transfection, activate Wnt/β-catenin signaling (**Fig. 5**).

Our transcriptomic studies comparing the mRNA expression profile of DDX5^KD^ HepAD38 cells, as a function of HBV replication and sorafenib treatment, further support the role of DDX5 as an upstream regulator of Wnt pathway activation, by regulating Wnt receptor expression. *FZD7* overexpression occurs in over 90% of chronic HCC patients (37) and dysregulation of Wnt/FZD receptors is common in HCC (50). Our RNA Seq analysis identified deregulated expression of several *FZD* receptors, co-receptors as well as regulators of Wnt signaling, upon DDX5 knockdown (**Figs. 5** and **Fig. 6**). Specifically, the deregulated expression of Wnt pathway genes *FZD7, SFRP4, SFRP5, WNT7B, MMP7* found in our study agrees with similarly deregulated expression of *FZD7* and *SFRP5* observed in HBV-related HCCs (50). Likewise, HBV-related liver tumors with reduced *DDX5* mRNA (7) exhibit increased expression of *FZD7* and *MMP7* mRNA (**Fig. 7B**). Lastly, the increased expression of Wnt receptors *FZD3, 4* and 7 in DDX5^KD^ cells, irrespective of HBV replication status, suggests that DDX5 is an upstream transcriptional regulator of Wnt pathway activation. Based on the well-established role of Wnt pathway activation in cellular reprogramming and pluripotency (33), our studies provide a mechanistic link of DDX5 loss to hepatocyte reprogramming/stemness in HCC.

Similar to decreased *DDX5* mRNA level, elevated expression of *FZD7* and *MMP7* is associated with poor overal survival in viral as well as non-viral associated HCCs (**Fig. 7**). Interestingly, our results also show that sorafenib treatment of HBV replicating DDX5^KD^ cells suppressed expression of the negative Wnt effectors *SFRP4* and *SFRP5*, while increasing expression of DVL1 and DVL3 (**Fig. 7**) which prevent β-catenin degradation (38). Collectively, our observations support that downregulation of DDX5 is a clinically relevant player in the pathogenesis of poor prognosis HBV-associated HCCs.

The mechanism by which DDX5 suppresses HBV biosynthesis requires further study. DDX5 could regulate viral transcription from cccDNA and/or translation from viral transcripts. Similarly, how DDX5 regulates stemness is incompletely understood. DDX5 was shown to act as a roadblock of somatic cell reprogramming by processing miR-125b, which in turn represses RING1 and YY1 Binding Protein (RYBP), a known inducer of pluripotency-associated genes (31). We also reported earlier that DDX5 is a positive regulator of PRC2 stability (7), and loss of PRC2 function during HBV replication leads to activation of Wnt signaling and re-expression of a hCSC-like gene signature (14). Hence, restoring DDX5 expression in chronically infected hepatocytes could suppress re-expression of the hCSC-like phenotype, providing therapeutic benefit.

Considering this important role of DDX5, we developed antagomirs against miR106b∼25 and miR17∼92 to restore DDX5 levels. Significantly, these antagomirs exert both antiviral effects reducing expression of HBV pgRNA and HBc, as well as anti-tumor effects restoring tumor suppressor PTEN and LKB1 (**Fig. 8**). Additionally, antagomir-mediated rescue of DDX5 reversed Wnt/β-catenin activation in HBV replicating cells, determined by loss of nuclear localization of β-catenin (**Fig. 8**). The results reported herein suggest antagomirs against miR106b∼25 and miR17∼92 can be explored as therapeutic strategy to suppress DDX5 downregulation. This strategy targets multiple pathways important for HCC, namely, (i) inhibition of HBV replication/biosynthesis (ii) rescue of tumor suppressor genes, and (iii) repression of Wnt signaling.

## Acknowledgments

The authors thank the French National Biological Resources Centre for frozen human liver tissues, obtained following approved consent from the French Liver Tumor Network Scientific Committee. The French Liver Tumor Network is funded by the Institut National de la Santé et de la Recherche Médicale (INSERM) and the Agence Nationale de la Recherche (ANR). The authors also thank Dr. Adam Zlotnick for generously providing the HBc antibody, and Dr. RL Hullinger for manuscript editing and image preparation.

