## Supporting Information for "RNA Helicase DDX5 Negatively Regulates Wnt Signaling and Hepatocyte Reprogramming in Hepatitis B Virus-related Hepatocellular Carcinoma"

and

Ourania Andrisani[1, 4#]

[1]Department of Basic Medical Sciences, [2]Department of Biochemistry, [3]Department of computer Science and [4]Purdue Center for Cancer Research, Purdue University, West Lafayette, IN 47907, [5]Cancer Research Center of Lyon UMR Inserm 1052 - CNRS 5286, and [6]Department of Hepatology, Hôpital de la Croix-Rousse, Hospices Civils de Lyon, Université Lyon 1, Lyon, France

*equal contribution

**Supporting data, Figure legends**

**Figure S1:** 3'UTR of DDX5 illustrating the complementarity of the seed sequence for each of the indicated microRNAs.

**Figure S2: (A)** RT PCR quantification of indicated miRNAs in human HBV-related HCC samples in comparison to peritumor. Tumor samples are classified into Group III *vs.* Rest (14), and DDX5 $^{high}$ *vs*. DDX5 $^{low}$ (7). **(B)**. RT PCR quantification of combined miR17~92 (mir-17+miR-20a) and miR-106b~25 (miR-106b+miR-93) in indicated HBV-related HCC samples. * $p<0.05$.

**Figure S3:** Dose response curves for cisplatin and sorafenib using HepAD38 cells, treated with the indicated concentrations for 2 days, n=3.

**Figure S4: (A)** Real time PCR analysis of miRNA expression in WT and stable miR17~92 and miR106b~25 overexpressing HepAD38 cells (miR$^{O/E}$). **(B)** MTS assay measuring proliferation of WT and stable miR17~92 and miR106b~25 overexpressing HepAD38 cells without or with HBV replication

**Figure S5:** Survival analysis (Km plots) of *MMP7*, *SFRP4*, *DVL1* and *DVL3* expression in 29 HCC patients who received sorafenib treatment.

**Figure S6**: **(A)** Immunoblots of indicated proteins without (-) or with (+) HBV replication for 5 days, in WT and DDX5$^{KD}$ HepAD38 cells. **(B)** Quantification of immunoblot data from three independent biological replicates for the indicated proteins from Fig. 8A. **(C)** Immunofluorescence staining of HBc in HepAD38 cells with HBV replication for 5 days. Negative control inhibitor or inhibitor cocktail containing miRNA inhibitors against miR-106b, miR-17, miR-20a and miR-19b1 was transfected on day 3 of HBV replication. (Right Panel) Quantification of percentage of cells showing HBc staining from 3 independent biological replicates. * $p<0.05$; Error bars indicate Mean ± SEM. **(D)** Quantification by ImageJ software of fluorescence intensities of HepAD38 cells transfected with fluorescent antagomir miR-19b1i-FAM (50 nM) or miR-17i-FAM (50 nM). Antagomirs were transfected on day4 of HBV replication and cells were fixed and immunostained on day 5 p.i. * $p<0.05$; Error bars indicate Mean ± SEM. **(E)** Immunofluorescence staining of HBc in HepG2-NTCP cells infected with 100 genome equivalents of HBV per cell. FAM conjugated miRNA inhibitor against miR-19b1 or miR-17 (green) was transfected on day 6 post infection.

### Supporting Materials and Methods

**Immunoblotting:** performed as previously described (12). Cells were lysed (15 mins, 4$^{o}$C) in lysis buffer (Cell Signaling), sonicated on ice for 30 secs and clarified by centrifugation (13,000 rpm, 15 mins, 4$^{o}$C). Protein concentration was determined using BCA assay. All samples were diluted to 1ug/ul using 4x dye (Biorad) and equal amounts of proteins (5µg - 40µg per lane) were run on SDS-PAGE. Following electrophoresis, proteins were transferred to nitrocellulose membrane via wet transfer (200mA, 45 - 90 mins at 4$^{o}$C). Following transfer, membranes were blocked with 3% (w/v) BSA in Tris-buffered saline containing 0.1% (v/v) Tween 20 (TBST) and incubated with primary antibody in 3% (w/v) BSA in TBST for 1hr at room temperature, followed by incubation in secondary antibody (1:2000 dilution) in 3% BSA in TBST for 1hr at room temperature. Three washes were performed after primary and secondary antibody incubations. Protein bands were detected by chemiluminescence using Pierce ECL (Biorad). Densitometric analysis of immunoblots was performed using ImageJ. Immunoblots are representative of three independent experiments. Antibodies used are listed in Supplementary Table S2:

**Immunofluorescence microscopy:** was performed as described (12). Cells (1x10$^{5}$) were seeded on coverslips in 12-well plates and incubated at 37$^{o}$C overnight. miRNA inhibitors were transfected using RNAimax and Lipofectamine 3000 on day 3. On day 5, cells were fixed using 4% PFA for 10 mins at room temperature. Following fixation, cells were permeabilized with Phosphate Buffered Saline (PBS) containing 0.1% (v/v) Triton X-100 for 15 mins at room temperature. Blocking was done using 10% goat serum in PBS containing 0.1% (v/v) Tween-20 (PBST) for 1 hour at room temperature. Primary antibodies diluted in blocking solution were used to detect HBc or beta-catenin for 1 hr at room temperature. Secondary antibodies conjugated to fluorophores were diluted in blocking solution and incubated for 45 mins at room temperature.

Three washes (5 min. each) were performed after primary and secondary antibody incubations. A 1 min incubation with Hoechst diluted in PBST to a final concentration of 2µg/ml to stain nuclei. Coverslips with cells were inverted and mounted in Antifade (10µl) on a glass slide. Microscopy was performed using an Olympus Fluoview confocal microscope. Antibodies used are listed in supplementary Table S2.

**RNA preparation and qRT-PCR:** RNA was isolated using Purelink mRNA Mini kit (Life Technologies) or Direct Zol RNA miniprep kit (Zymo Research). cDNA synthesized from 1.0 µg total RNA using iSCRIPT cDNA synthesis kit (Biorad). qRT-PCR performed using SYBR green (Roche) in triplicates, normalized to GAPDH. For miRNA quantification, RNA was extracted using miRNeasy Mini Kit (Qiagen) or Trizol for liver tissues from HBV patients with HCC (tumor and peritumoral tissue). Liver tumor samples were obtained from the French National Biological Resources Centre following approved consent from the French Liver Tumor Network Scientific Committee. cDNA synthesized from 1.0 µg RNA using HiSpec buffer in miScript II RT Kit (Qiagen), and qRT-PCR performed using miScript SYBR® Green PCR Kit (Qiagen).

**Supporting Table S1: List of Plasmids and siRNAs**

| Plasmids, siRNAs | Source |
|---|---|
| TOPFlash vector[(1)] | Addgene (12456) |
| FOPFlash vector[(1)] | Addgene (12457) |
| Renilla luciferase vector[(3)] | Addgene (27163) |
| Empty vector[(2)] | (43) |
| miR106b~25 vector[(2)] | (43) |
| miR17~92 vector[(4)] | Addgene (21109) |
| Firefly luciferase - 3'UTR DDX5 in pMirTarget vector | Origene (#SC215943) |
| Ctrlsi | ThermoFisher Scientific (#4390843) |
| DDX5si#1 | ThermoFisher Scientific (#4392420, assay id s4007) |
| DDX5si#2 | ThermoFisher Scientific (#4392420, assay id s4008) |
| hsa-miR-19b-1-5p miRCURY LNA miRNA Power Inhibitor | Qiagen (Product# 339131, Catalog# YI04100832-DDB) |
| hsa-miR-17-5p miRCURY LNA miRNA Power Inhibitor | Qiagen (Product#339131; Catalog# YI04100215-DDB) |


(1) Veeman MT, Slusarski DC, Kaykas A, Louie SH, Moon RT. (2003). Zebrafish prickle, a modulator of noncanonical Wnt/Fz signaling, regulates gastrulation movements. Curr Biol.; 13(8):680-5.

(2) Smith, A.L., Iwanaga, R., Drasin, D.J., Micalizzi, D.S., Vartuli, R.L., Tan, A.C., and Ford, H.L. (2012). The miR-106b-25 cluster targets Smad7, activates TGF-beta signaling, and induces EMT and tumor initiating cell characteristics downstream of Six1 in human breast cancer. Oncogene. 31, 5162-5171.

(3) Chen X, Prywes R. (1999). Serum-induced expression of the cdc25A gene by relief of E2F-mediated repression. Mol Cell Biol. 19(7):4695-702.

(4) O'Donnell KA, Wentzel EA, Zeller KI, Dang CV, Mendell JT. (2005). c-Myc-regulated microRNAs modulate E2F1 expression. Nature. 435(7043):839-43.

**Supporting Table S2: Antibodies Used**

| Antibody | Dilution | Application | Source |
|---|---|---|---|
| Rabbit α-Human DDX5 | 1:1000 in 3% BSA in TBST | Western Blot | Cell Signaling Technologies (#9877S) |
| Mouse α-Human Actin | 1:2000 in 3% BSA in TBST | Western Blot | Sigma (#A5441) |
| Rabbit α-Human MCM7 | 1:1000 in 3% BSA in TBST | Western Blot | Cell Signaling Technologies (#3735S) |
| Rabbit α-Human OCT4 | 1:1000 in 3% BSA in TBST | Western Blot | Cell Signaling Technologies (#2750S), Abcam (#ab19857) |
| Mouse α-Human SOX2 | 1:1000 in 3% BSA in TBST | Western Blot | Cell Signaling Technologies (#4900S) |
| Rabbit α-Human NANOG | 1:1000 in 3% BSA in TBST | Western Blot | Cell Signaling Technologies (#4903S) |
| Mouse α-Human CD44 | 1:1000 in 3% BSA in TBST | Western Blot | Cell Signaling Technologies (#3570S) |
| Rabbit α-Human PTEN | 1:1000 in 3% BSA in TBST | Western Blot | Cell Signaling Technologies (#9559S), R&D Systems (# AF847) |
| Rabbit α-Human LKB1 | 1:1000 in 3% BSA in TBST | Western Blot | Cell Signaling Technologies (#3047S), Novus Biologicals (#NBP2-14835SS) |
| Rabbit α-Human Akt | 1:1000 in 3% BSA in TBST | Western Blot | Cell Signaling Technologies (#4691S) |
| Rabbit α-Human p-Akt | 1:1000 in 3% BSA in TBST | Western Blot | Cell Signaling Technologies (#4060S) |
| Mouse α-Human SMAD7 | 1:1000 in 3% BSA in TBST | Western Blot | R&D Systems (# MAB2029) |
| Rabbit α-HBV Core | 1:5000 in 3% BSA in TBST | Western Blot | Dr.Adam Zlotnick lab |

| | | | |
|---|---|---|---|
| Horse α-Mouse secondary | 1:2000 in 3% BSA in TBST | Western Blot | Vector Laboratories (#PI-2000) |
| Goat α-Rabbit secondary | 1:2000 in 3% BSA in TBST | Western Blot | Vector Laboratories (#PI-1000) |
| Rabbit IgG | 5μg | ChIP | Cell Signaling Technologies (#2729S) |
| Rabbit α-Human c-Myc | 5μg | ChIP | Cell Signaling Technologies (#9402S) |
| Rabbit α-Human DDX5 | 1:100 in 10% Goat serum in PBST | IF | Cell Signaling Technologies (#9877S) |
| Mouse α-Human β-catenin | 1:100 in 10% Goat serum in PBST | IF | Cell Signaling Technologies (#2677S) |
| Rabbit α-HBV Core | 1:1000 in 10% Goat serum in PBST | IF | Dr. Adam Zlotnick |
| Goat α-Rabbit Alexa Fluor 633 | 1:2000 in 10% Goat serum in PBST | IF | ThermoFisher Scientific (#A21070) |
| Goat α-Rabbit Alexa Fluor 594 | 1:2000 in 10% Goat serum in PBST | IF | ThermoFisher Scientific (#A11012) |
| Goat α-Rabbit Alexa Fluor 488 | 1:2000 in 10% Goat serum in PBST | IF | ThermoFisher Scientific (#A11008) |
| Goat α-Mouse Alexa Fluor 633 | 1:2000 in 10% Goat serum in PBST | IF | ThermoFisher Scientific (#A21050) |

**Supporting Table S3: Primer sequences**

| Primer | 5’ - Sequence - 3’ |
|---|---|
| DDX5 Forward | AGCAAGTGAGCGACCTTATC |
| DDX5 Reverse | CATCCTTCATGCCTCCTCTAC |
| EpCAM Forward | TCGTCAATGCCAGTGTACTTC |
| EpCAM Reverse | GCCATTCATTTCTGCCTTCATC |
| AFP Forward | AGACTGAAAACCCTCTTGAATGC |
| AFP Reverse | GTCCTCACTGAGTTGGCAACA |
| OCT4 Forward | GTG TTC AGC CAA AAG ACC ATC T |
| OCT4 Reverse | GGC CTG CAT GAG GGT TTC T |
| SOX2 Forward | TGG ACA GTT ACG CGC ACA T |
| SOX2 Reverse | CGA GTA GGA CAT GCT GTA GGT |
| NANOG Forward | TTT GTG GGC CTG AAG AAA ACT |
| NANOG Reverse | AGG GCT GTC CTG AAT AAG CAG |
| FZD7 Forward | GCCTGATGTACTTTAAGGAGGAG |
| FZD7 Reverse | CAGGTAGGTGAGAACGGTAAAG |
| WNT7B Forward | TCTACGTGTTTCTCTGCTTTGG |
| WNT7B Reverse | GCTAGGCCAGGAATCTTGTT |
| SFRP4 Forward | GCCAACTTTGGCAACGTATC |
| SFRP4 Reverse | CCACCGTTGTGACCTCATT |
| SFRP5 Forward | GATGTGCTCCAGTGACTTTGT |
| SFRP5 Reverse | GGCTTGAGCAGCTTCTTCTT |
| DVL1 Forward | GACTCATCCGGAAGCACAAA |
| DVL1 Reverse | GACATGGTGGAGTCGGTTATG |

| | |
|---|---|
| DVL3 Forward | CGGCATCTACATTGGCTCTATC |
| DVL3 Reverse | CGGACTGCATCGTCATTACTC |
| MMP7 Forward | GCTCACTTCGATGAGGATGAA |
| MMP7 Reverse | AGGAATGTCCCATACCCAAAG |
| GAPDH Forward | CCCTTCATTGACCTCAACTACA |
| GAPDH Reverse | ATGACAAGCTTCCCGTTCTC |
| HBc Forward | CTGGGTGGGTGTTAATTTGG |
| HBc Reverse | TAGGGGCATTTGGTGGTCTA |
| HBV pgRNA Forward | CTCCTCCAGCTTATAGACC |
| HBV pgRNA Reverse | GTGAGTGGGCCTACAAA |
| Total HBV RNA Forward | TCACCAGCACCATGCAAC |
| Total HBV RNA Reverse | AAGCCACCCAAGGCACAG |
| HBV pgRNA (Protzer) Forward | GAGTGTGGATTCGCACTCC |
| HBV pgRNA (Protzer) Reverse | GAGGCGAGGGAGTTCTTCT |
| miR-17 5p Forward | CAAAGTGCTTACAGTGCAGGTAG |
| miR-18a 5p Forward | TAAGGTGCATCTAGTGCAGATAG |
| miR-19a 5p Forward | TGTGCAAATCTATGCAAAACTGA |
| miR-19b1 5p Forward | TGTGCAAATCCATGCAAAACTGA |
| miR-20a 5p Forward | TAAAGTGCTTATAGTGCAGGTAG |
| miR-92a1 5p Forward | TATTGCACTTGTCCCGGCCTGT |
| miR-106b 5p Forward | TAAAGTGCTGACAGTGCAGAT |
| miR-93 5p Forward | CAAAGTGCTGTTCGTGCAGGTAG |
| miR-25 5p Forward | CATTGCACTTGTCTCGGTCTGA |
| U6 snRNA Forward | CTCGCTTCGGCAGCACATATACT |

| | |
|---|---|
| U6 snRNA Reverse | ACGCTTCACGAATTTGCGTGTC |
| Universal Reverse | Qiagen Propriety information |
| c-myc site 1 Forward | ACCTCGGAAACCCACCAAG |
| c-myc site 1 Reverse | TCTCCCTGGGACTCGACG |
| c-myc site 2 Forward | AAAGGCAGGCTCGTCGTTG |
| c-myc site 2 Reverse | CGGGATAAAGAGTTGTTTCTCCAA |
| c-myc site 3 Forward | CTCGACTCTTACTCTCACAAATGG |
| c-myc site 3 Reverse | GCTACTGGTGCAGTTAGGTCC |
| miR-106b 5p inhibitor | mA/ZEN/mU mCmUmG mCmAmC mUmGmU mCmAmG mCmAmC mUmUmU /3ZEN/ |
| miR-17 5p inhibitor | mC/ZEN/mU mAmCmC mUmGmC mAmCmU mGmUmA mAmGmC mAmCmU mUmU/3ZEN/ |
| miR-19b1 inhibitor | mG/ZEN/mC mUmGmG mAmUmG mCmAmA mAmCmC mUmGmC mAmAmA mAmC/3ZEN/ |
| miR-20a inhibitor | mC/ZEN/mU mAmCmC mUmGmC mAmCmU mAmUmA mAmGmC mAmCmU mUmU/3ZEN/ |
| Negative Control inhibitor | mG/ZEN/mC mGmAmC mUmAmU mAmCmG mCmGmC mAmAmU mAmUmG mG/3ZEN/ |

**Supporting Table S4: Reagents, Chemical inhibitors, and Kits**

| **Reagents, Kits** | **Source** |
|---|---|
| Cisplatin | Selleck Chemicals (#S1166) |
| Sorafenib | Selleck Chemicals (#S7397) |
| XAV-939 | Selleck Chemicals (#S1180) |
| ICG-001 | Selleck Chemicals (#S2662) |
| Dual-Luciferase® Reporter Assay System | Promega (#E1980) |

| | |
|---|---|
| CellTiter 96® AQueous One Solution Cell Proliferation Assay (MTS) | Promega (#G3580) |
| Cell Lysis Buffer (10X) | Cell Signaling Technology (#9803) |
| miScript SYBR® Green PCR Kit | Qiagen (#218073) |
| miScript II RT Kit | Qiagen (#218161) |
| miRNeasy Mini Kit | Qiagen (#217004) |
| LightCycler® 480 SYBR Green I Master | Roche (#04887352001) |
| LightCycler® 8-Tube Strips (white) | Roche (#06612601001) |
| LightCycler® 480 Sealing Foil | Roche (#04729757001) |
| 4x Laemmli Sample Buffer | Biorad (#1610747) |
| iScript™ cDNA Synthesis Kit | Biorad (#1708891) |
| Nitrocellulose Membrane, Roll, 0.2 µm | Biorad (#1620112) |
| DMSO | Sigma (#D8418-50ML) |
| Tetracycline hydrochloride | Sigma (#T7660-5G) |
| Bovine Serum Albumin | Sigma (#A9647-100G) |
| Triton™ X-100 | Sigma (#T8787-100ML) |
| Costar® 6-well Ultra-Low Attachment Plates | Corning (#3471) |
| Pierce™ ECL Western Blotting Substrate | ThermoFisher Scientific (#32106) |
| Geneticin™ Selective Antibiotic (G418 Sulfate) | ThermoFisher Scientific (#10131027) |
| PureLink™ RNA Mini Kit | ThermoFisher Scientific (#12183025) |
| Pierce™ BCA Protein Assay Kit | ThermoFisher Scientific (#23227) |
| Lipofectamine™ 3000 Transfection Reagent | ThermoFisher Scientific (#L3000015) |
| Lipofectamine™ RNAiMAX Transfection Reagent | ThermoFisher Scientific (#13778150) |
| Hoechst 33342 Solution (20 mM) | ThermoFisher Scientific (#62249) |
| Tween™ 20 | ThermoFisher Scientific (BP337-500) |

| | |
|---|---|
| ProLong™ Diamond Antifade Mountant | ThermoFisher Scientific (#P36970) |
| B-27™ Supplement (50X), minus vitamin A | ThermoFisher Scientific (#12587010) |
| Recombinant Human EGF | Peprotech (#AF-100-15) |
| Recombinant Human FGF-basic | Peprotech (#100-18C) |
