## Supporting Figs 1 to 6 for "RNA Helicase DDX5 Negatively Regulates Wnt Signaling and Hepatocyte Reprogramming in Hepatitis B Virus-related Hepatocellular Carcinoma"

**A**

```
DDX5 3'UTR (129-135)    5' GCACUUU   3'
                             ||||||
miR17, 20a, 106b, 93    3'  GUGAAA   5'

DDX5 3'UTR (129-135)    5' GCACUUU   3'
                             |||  ||
miR18a                  3'  GUGGAA   5'

DDX5 3'UTR (129-135)    5' GCACUUU   3'
                                ||||
miR92-1, 25             3'  ACGUUA   5'

DDX5 3'UTR (129-135)    5' GCACUUU   3'
                             |||  ||
miR19a, 19b-1           3'  GUGCAA   5'
```

Fig. S1 Mani et al.

A

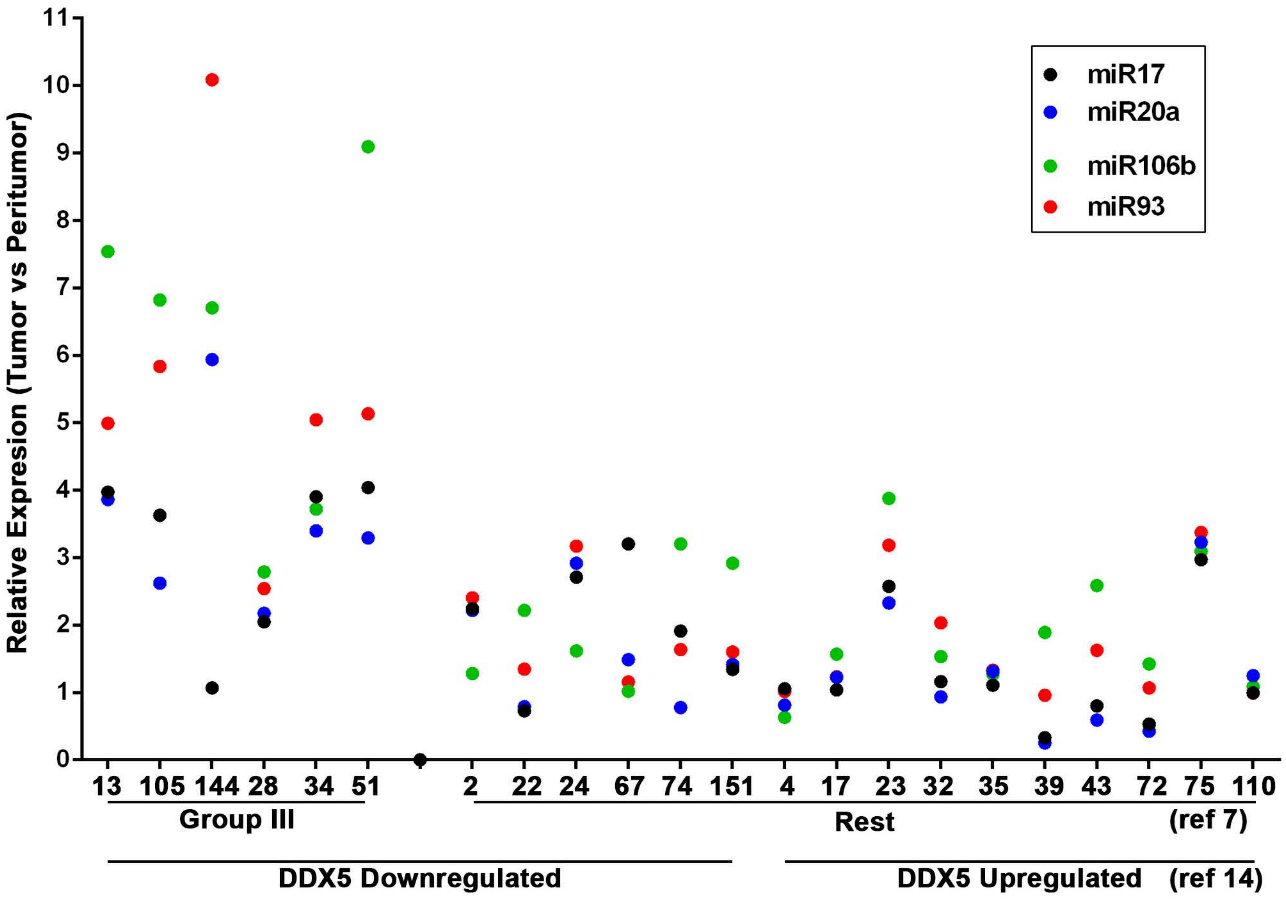


B

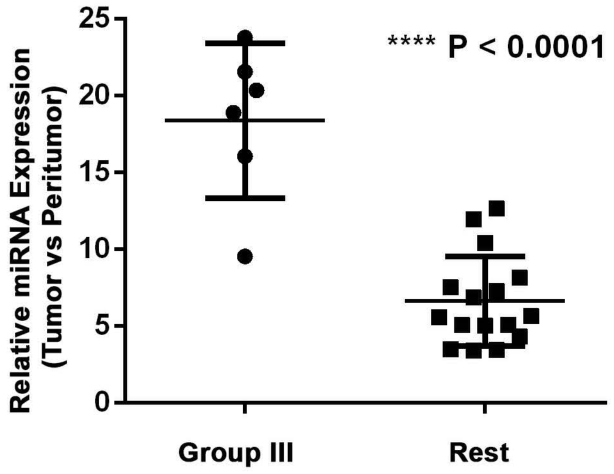


Fig. S2 Mani et al.

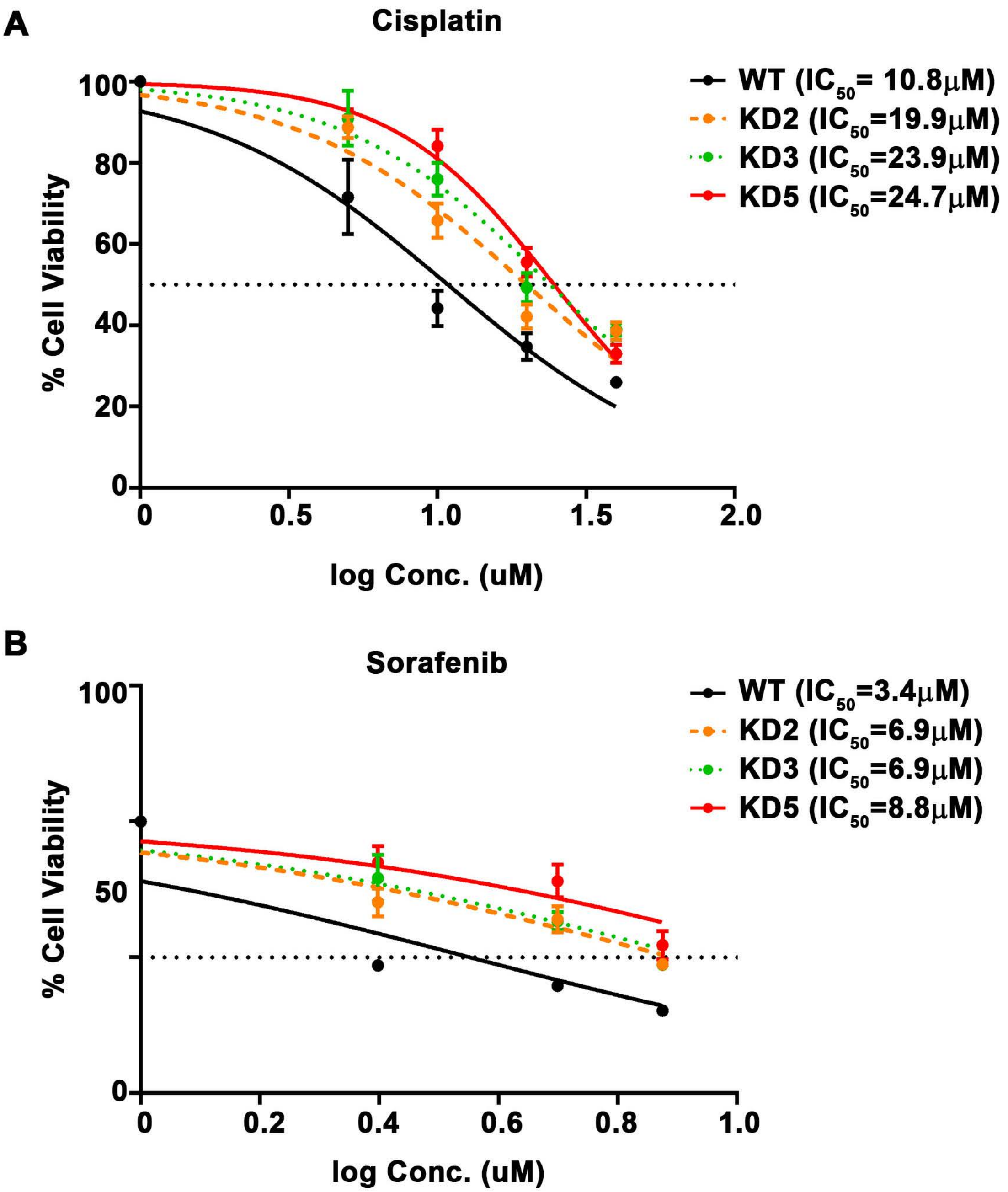


Fig. S3 Mani et al.

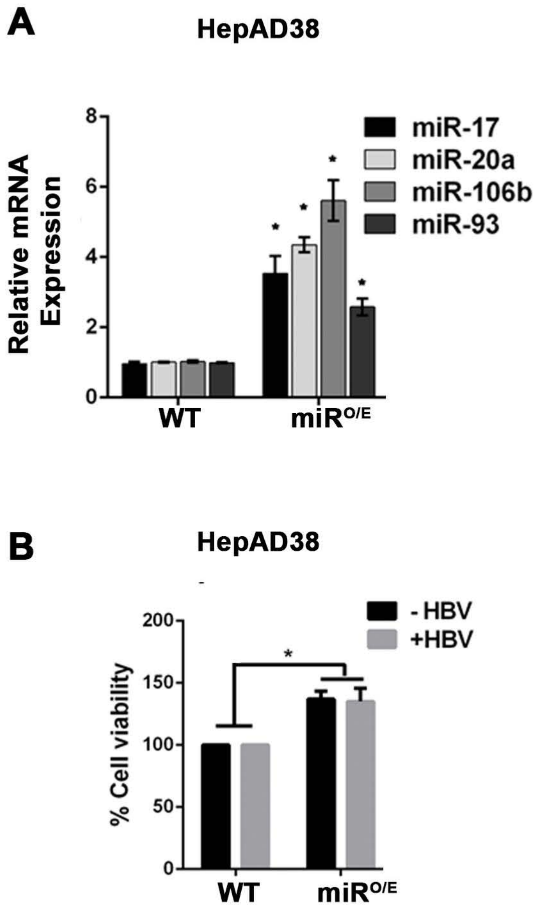


**Fig. S4 Mani et al.**

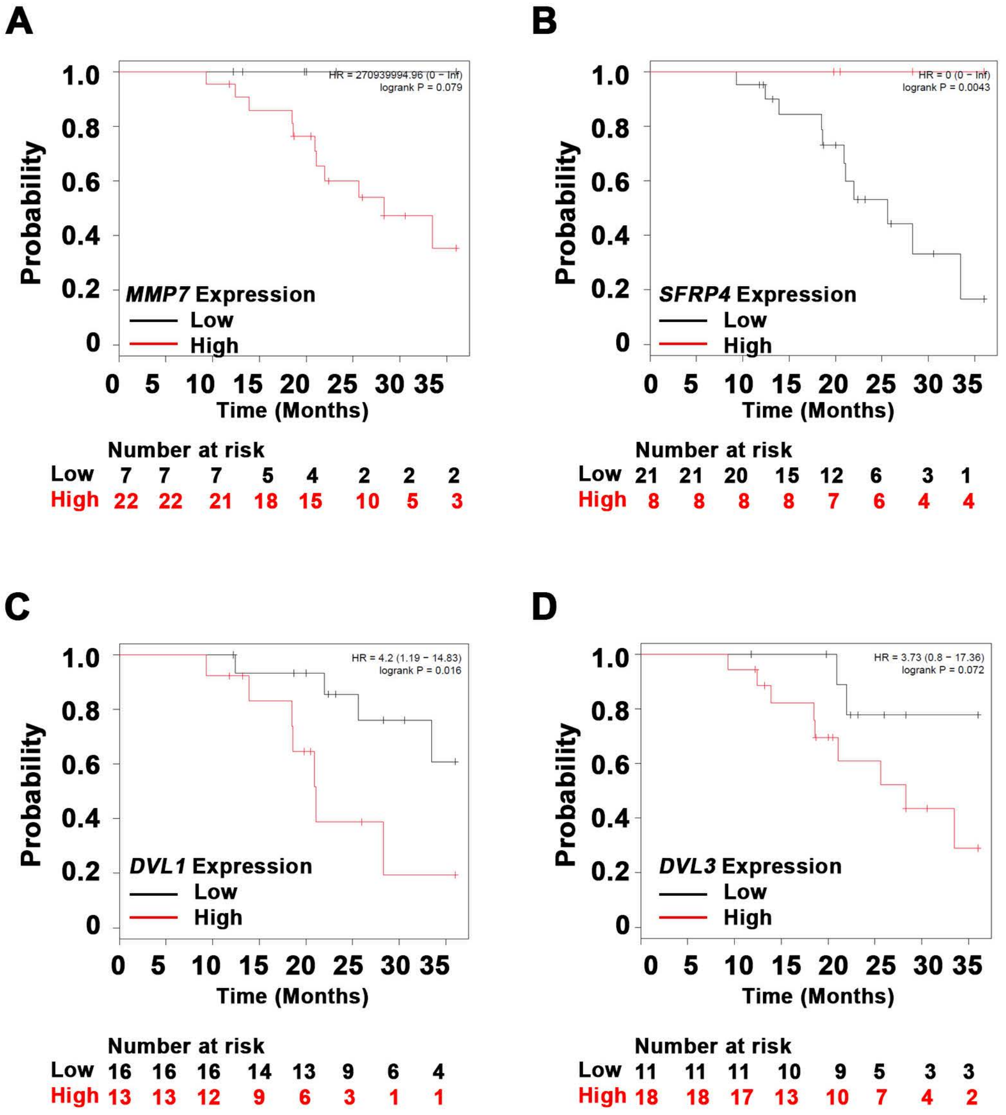


**Fig. S5 Mani et al.**

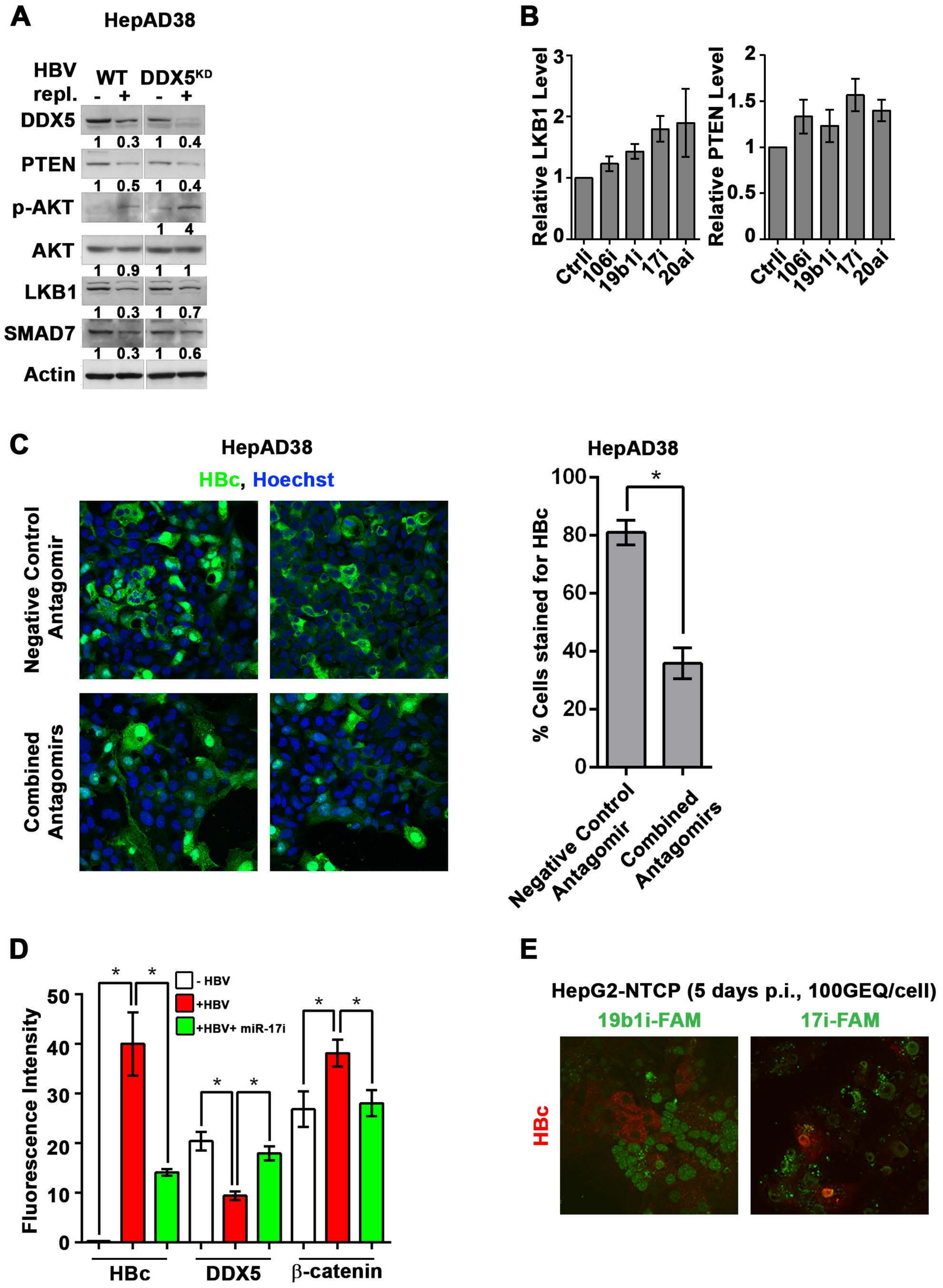


Fig. S6 Mani et al.
